# Amyloid signaling in antiphage defense

**DOI:** 10.1101/2024.11.07.622273

**Authors:** Léa Ibarlosa, Sonia Dheur, Corinne Sanchez, Seamoon Deb, Alexandra Granger-Farbos, Virginie Coustou, Mélanie Berbon, Aurélie Massoni, Bénédicte Salin, Corinne Blancard, Claire Saragaglia, Xavière Menatong Tene, Nicolas Dufour, Ombeline Rossier, Birgit Habenstein, Laurent Debarbieux, Brice Kauffmann, Antoine Loquet, Sven J. Saupe

## Abstract

Immune regulated cell death (RCD) is a defense strategy common to different domains of life involving purposeful sacrifice of infected cells. In animals and fungi, functional amyloids play a role in the control of RCD as molecular switches activating key cell death effectors, rather than causing direct toxicity as seen with pathological amyloids. Here, we describe a novel amyloid-based signal transduction mechanism in an antiphage defense system in *Escherichia coli*. This antiphage abortive infection (Abi) system is mediated by two proteins, Bab and Agp which share a common amyloid motif and are encoded by adjacent genes. Upon phage infection, Agp activates Bab through amyloid signaling, leading to cell death. We determined the structure of the Bab cell death execution domain, which is distantly related to pore-forming domains present in fungi, animals and plants. We show that Bab and the fungal HET-S amyloid-controlled cell death execution protein are functionally interchangeable in their respective roles in antiphage defense and allorecognition. These findings add antiphage defense to the functional repertoire of amyloids.

## Introduction

Hundreds of defense systems enabling bacteria to combat phage infection have been identified, with many having counterparts in human cell-autonomous immune pathways ^1-3^. Antiphage systems are broadly classified into two categories: those that directly interfere with phage replication and Abortive Infection (Abi) systems that trigger death of infected host cells ^4^. Abi defense represents a form of regulated cell death (RCD), a widespread immune strategy common to different domains of life ^5^. Amyloid cross-β protein assemblies, often associated with neurodegenerative diseases ^6^, have been found to ensure diverse functional roles, including the regulation of RCD in animals and fungi ^7, 8^. In mammals, RHIM amyloid motifs are crucial for the formation of the RIP1/RIP3 kinase complex that controls necroptosis and in Drosophila, cRHIM motifs mediate both antibacterial and antiviral defenses ^9-11^. As counter-defense, viral and bacterial pathogens utilize RHIM motifs to disrupt immune responses^8^. In fungi, in a process termed amyloid signaling, Nod-like receptors (NLRs) activate their cognate downstream cell death effector proteins by converting short regulatory motifs to an amyloid cross-β conformation ^12-15^. HET-S of *Podospora anserina*, the paradigm of such fungal amyloid-controlled cell death effectors, is involved in heterokaryon incompatibility (HI), an RCD reaction occurring when strains displaying different incompatibility determinants fuse. HI is a non-self recognition process, which prevents cytoplasmic exchanges between genetically unlike strains and therefore limits mycovirus transmission ^12, 16^ . HET-S contains a pore-forming domain (designated HeLo) and a C-terminal amyloid motif ^17, 18^. Incompatibility is initiated when HET-S interacts with the alternate incompatibility determinant, the [Het-s] prion amyloid protein. [Het-s] acts as a template, converting the amyloid motif of HET-S, which subsequently induces refolding and activation of the HeLo domain^13^. HET-S is also the cognate downstream cell death effector of NWD2, a NACHT domain NLR displaying an N-terminal amyloid motif (termed R0). NWD2 is encoded by a gene immediately adjacent to *het-S*^19^. Ligand induced oligomerisation of NWD2 leads to amyloid conversion of the R0 motif, which likewise activates HET-S. Several additional fungal amyloid signaling sequences (FASS) have been identified including a motif similar to RHIM (designated PP) ^14, 20, 21^. FASS motifs reside in products of adjacent gene pairs associating NLRs to HeLo or Hell (HeLo-like) cell death effectors. FASS motifs (and RHIM) can function as prion-forming domains *in vivo* ^14, 15, 21, 22^. Similar adjacent gene pairs encoding proteins with shared amyloid prion motifs have been identified in multicellular bacteria ^23^. These bacterial amyloid signaling sequences (BASS) have been classified into ten families and among them is BASS3 which is similar to RHIM and PP ^21, 23^. BASS motifs, like their fungal counterparts, are located at the N-termini of NB-ARC and NACHT domain NLRs, and C-termini of Bell (a Hell-related domain) and other domains with immune functions such as TIR or CHAT. Hell and Bell domains belong to a broader family of membrane targeting domains with immune functions including the pore-forming domain of MLKL, which controls necroptosis and CC^HeLo^ of plant NLR resistosomes ^14, 23, 24,25, 26^. The amyloid forming properties of several BASS-motifs have been confirmed but the biological function of bacterial Bell/NLR gene pairs remains to be elucidated ^23^.

Comparative genomics approaches have identified a novel protein architecture suspected to be involved in antiphage defense ^27^. This architecture, referred to as TGP, consisting of TPR repeats, a GreA/B-C RNA polymerase-binding domain and a PIN nuclease domain, is often linked to defense-related domains such as TIR, EAD10, or PNPase ^27^. Kaur and colleagues hypothesize that during phage infection, TGP proteins bind to RNA polymerase hijacked for viral transcription and degrade the corresponding transcripts via their PIN domain. It is further proposed that if this first line of defense fails, TGP proteins activate associated downstream Abi-effector modules ^27^. Building on this model, we analyze here Bab/Agp, a gene pair from *E. coli* ECOR25 associating a Bell-domain protein to a TGP protein. We find that Bab and Agp share an amyloid motif we term BASS11 residing at the C-terminus of Bab and N-terminus of Agp. We establish that Bab/Agp confers resistance to phage T5 and other *Siphoviridae*. We find that T5 infection triggers Agp-dependent conversion of the amyloid motif of Bab and activates its globular domain to cause Abi. We present the structure of the Bab cell death execution domain. We redesign Bab into an artificial incompatibility system and HET-S into an antiphage system, underscoring the mechanistic resemblance of amyloid-controlled bacterial Abi and fungal RCD.

### Bab and Agp display an amyloid motif

In *E. coli* strain ECOR25, a gene encoding a TPR-GreA/B-C-PIN protein (we term Agp) is immediately adjacent to a gene encoding Bab, a protein with a small helical domain related to Bell-domains ^23, 27^(Fig. 1A, extended data Fig. 1). The N-terminal region of Bab (∼15 amino acids) is conserved in Hell and Bell domains and NLRs involved in antiphage defense (Avast2, bNACHT02, bNACHT11 and bNACHT23) ^15, 21, 23, 28, 29^ (extended data Fig. 1A). Agp and Bab display respectively at their N- and C-terminus, a predicted amyloid forming region (Fig.1A, extended data Fig. 1A and F and Supplementary Table S2). This conserved motif, we term BASS11 is also associated to other immunity-related domains such as TIR, CHAT, EAD7, NACHT and NB-ARC and occurs in products of adjacent genes (extended data Fig. 1B). The Bab/Agp gene pair is located in a defense island in a prophage sequence (hotspot #11) ^30^. The same is true for three additional Bab/Agp homologs (located in hotspots #1, 8 and 12), (extended data Fig. 2).

**Figure 1.**
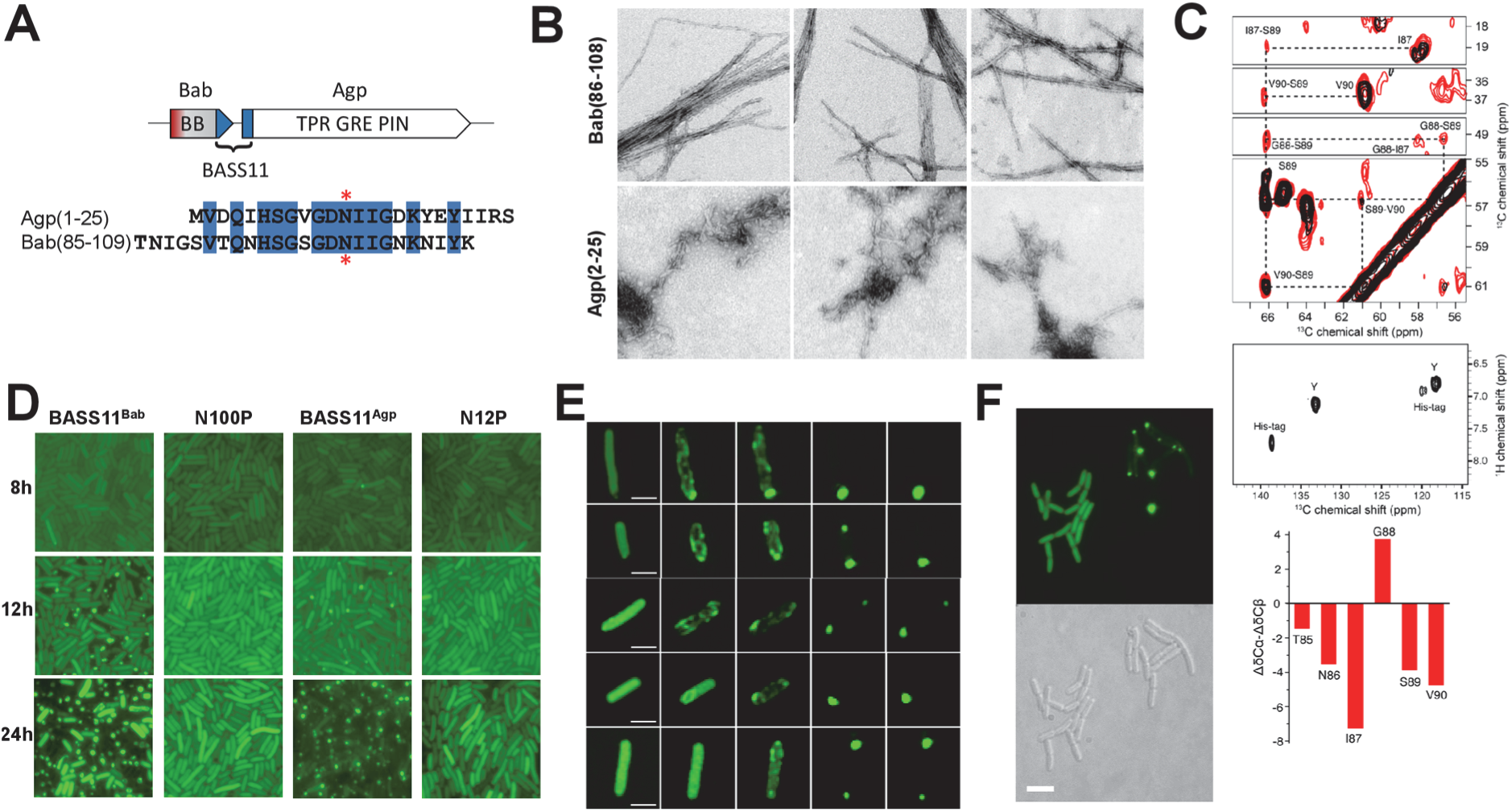
Bab and Agp share an amyloid forming motif with prion propertiesA. Domain architecture diagram of the Bab/Agp gene pair. Bab (OWC60847.1) encodes a 109 amino acid long protein with an N-terminal helical domain (BB for Babybell) with a predicted N-terminal hydrophobic α-helix (given in red) and a C-terminal predicted amyloid forming motif (given in blue). Agp (OWC60848.1) encodes a 1094 amino acid long protein with a TPR domain, a GreA/B-C RNA polymerase interaction domain and a PIN nuclease domain. An alignment of the predicted amyloid forming motifs of Bab and Agp is given. Positions of Agp N12 and Bab N100 targeted for mutation are marked with an asterisk. **B.** Negative staining electron micrographs of fibrils formed by Bab(86-108) and Agp(2-25) synthetic peptides. Each image is a 500 x 500 nm panel. **C.** Structural characterization of Bab (50-109) fibrils by solid-state NMR spectroscopy. Top panel, excerpts of 2D ^13^C-^13^C correlation experiments recorded at 800 MHz 1H frequency using mixing times of 50 ms (in black) and 150 ms (in red). Sequential assignment of the stretch I87-G88-S89-V90 is highlighted. Middle panel, excerpt of the aromatic region of the 2D ^1^H-^13^C INEPT, detecting disordered residues. Bottom panel, secondary chemical shifts for the stretch T85-V90. Positive or negative values correspond to α-helical or β-strand conformation, respectively. **D.** Dot-formation in *E. coli* cells expressing either BASS11^Bab^ or BASS11^Agp^ fused to GFP or the corresponding N100P and N12P mutants. Cells were imaged at different times after transformation as given. Each image is a 20 x 20 µm panel. Quantification of dot-formation is given in extended data Fig.3D. **F.** Time lapse microscopy of dot-formation in five different cells, time between frames is 30 min, scale bar is 2 µ. **F.** Transmission of the dot and diffuse state of BASS11^Bab^-GFP in cell divisions leading to formation of microcolonies homogenously displaying cells with dot or diffuse-fluorescence. Further illustration and quantification of dot-colony formation are given in extended data Fig. 4C and D.

Synthetic peptides corresponding to the BASS11 motifs of Bab (residue 86 to 108, BASS11^Bab^) and Agp (residue 2 to 25, BASS11^Agp^) form fibrils *in vitro* (Fig. 1B). The same was true for purified full-length Bab (extended data Fig. 2A). BASS11^Bab^ and BASS11^Agp^ fibrils induce ThT fluorescence and bind Congo red (extended data Fig. 2B and C). The recombinant C-terminal region of Bab (Bab(50-109)) spontaneously polymerized into amyloid fibrils and the structural conformation was probed with magic-angle spinning solid-state NMR (ssNMR). Two-dimensional ^13^C-^13^C correlation experiments confirmed the predominance of a rigid and immobilized core formed by ∼ 25 residues in a β-strand conformation. In particular, the stretch from residue 85 to 90 presents ^13^C secondary chemical shifts ^31^ indicating a β-sheet structure for residues 85-87 and 89-90, separated by a putative kink induced by glycine 88 (Fig. 1C). Several amino acids were only observed in a two-dimensional ^1^H-^13^C INEPT that reflect disordered residues, as exemplified with the His-tag and tyrosine 57 (Fig. 1C), indicating that a part of the sequence is not involved in the rigid amyloid core. AlphaFold (AF) predicted a four β-strand amyloid fold for the Bab BASS11 region with 4 glycine residues acting as β-sheet breakers (extended data Fig. 1F), flanked by disordered N- and C-terminal residues, the model was consistent with chemical shifts experimentally determined by ssNMR.

Previously characterized FASS/BASS motifs are able to function as prion forming domains and direct formation of transmissible foci as GFP fusion proteins in fungal model systems ^14, 15, 21-23, 32^. In *E. coli,* amyloid prion formation and transmission was also followed using GFP fusion proteins ^33^. We thus determined whether BASS11 could direct formation of transmissible GFP foci in *E. coli*. Cells were transformed with a plasmid expressing BASS11^Agp^-GFP or BASS11^Bab^-GFP and GFP-foci formation was analyzed at various time points after transformation. Fluorescence was initially diffuse in all cells (Fig. 1D). The proportion of cells with foci then increased over time and reached 70-80%, 24h after transformation (Fig. 1D, extended data Fig. 3D). Transition to the foci state was associated with insolubility of the GFP fusion protein (extended data Fig.3E). Putative β-breaker mutations, based on the AF model were designed in a central conserved position of BASS11^Agp^ and BASS11^Bab^ and they abolished foci formation (respectively N12P and N100P) (Fig. 1D, extended data Fig. 3D). In time lap microscopy, the transition from diffuse to foci state occurs with an intermediate state with disperse mesh-like aggregates that then coalesce into a single dot or a pair of dots (Fig. 1E). GFP-dots colocalized with refringent bodies at the cell poles (extended data Fig. 4A). In a small fraction of cells (3.4±0.6%), cable-like structures were also observed, as previously described during prion amyloid motif expression in yeast or *E. coli* (extended data Fig. 4B) ^32, 34, 35^. To determine whether the foci state is transmissible, liquid cultures containing cells with diffuse fluorescence and cells with foci were plated onto solid medium. The resulting colonies were homogenously composed of cells with diffuse fluorescence or cells with foci (Fig. 1F, extended data Fig.4C). A proportion of chimeric colonies was also observed with the diffuse and foci states segregating into distinct sectors (extended data Fig.4C). The proportion of dot-colonies gradually increased with the proportion of dot-cells in the starting cultures (extended data Fig.4D). Formation of homogenous colonies with either diffuse fluorescence or foci was reported previously for amyloid prion motifs expressed in *E. coli* or yeast and results from stable transmission of the aggregated state in cell divisions ^32, 35, 36^.

### Bab/Agp is an Abi antiphage defense system

We set out to assay for a possible antiphage activity of Bab/Agp. Attempts to clone Bab/Agp on plasmid failed, suggesting toxicity. Accordingly, arabinose-inducible expression of Bab/Agp was lethal (extended data Fig.5A, B and C). Toxicity was abolished by β-breaker N12P or N100P mutations in Agp and Bab and when Agp and Bab were expressed independently. Yet, strong overexpression of Bab alone (but not Bab N100P) was also toxic (extended data Fig.5D, E and F).

In order to obtain cells expressing Bab/Agp at viable levels, we constructed a strain with a chromosomal insertion of the gene pair with its native promotor. Bab/Agp was PCR-amplified from genomic DNA of *E. coli* ECOR25 and inserted at the *galM/gpmA* locus in a MG1655 derivate (using the *tetA-sacB* gene replacement system developed by the Court lab ^37^) and the resulting strain was infected with a set of phylogenetically diverse phages (P1, T4, T5, T6 and T7). MG1655*::Bab/Agp* was sensitive to P1, T4, T6 and T7, but no plaque formation was observed after T5 infection (Fig.2A, extended data Fig. 6A). In spot assays, growth inhibition halos formed at high phage titers but discrete plaques were not observed (Fig.2B, extended data Fig.6A). Growth assays in liquid culture confirmed that the *Bab/Agp* strain is completely resistant to T5 at low MOI (0.01). At higher MOI, there was a growth arrest of both *Bab/Agp* and the control MG1655 strain (extended data Fig.6C). Similarly, in plating assays, at low MOI, viability in *Bab/Agp* was maintained but high MOI resulted in a loss of plating efficiency (extended data Fig.6B). Loss in plating efficiency was not associated to a loss of turbidity. In contrast to the MG1655 control, cell ghosts were present in T5 infected *Bab/Agp* cultures, thus growth arrest and reduction in plating efficiency do not result from viral-induced lysis (extended data Fig.6D-F). In cultures supernatants of infected *Bab/Agp*, there was no increase in phage titer further indicating that the strain is incompetent for phage replication (extended data Fig.6C).

**Figure 2.**
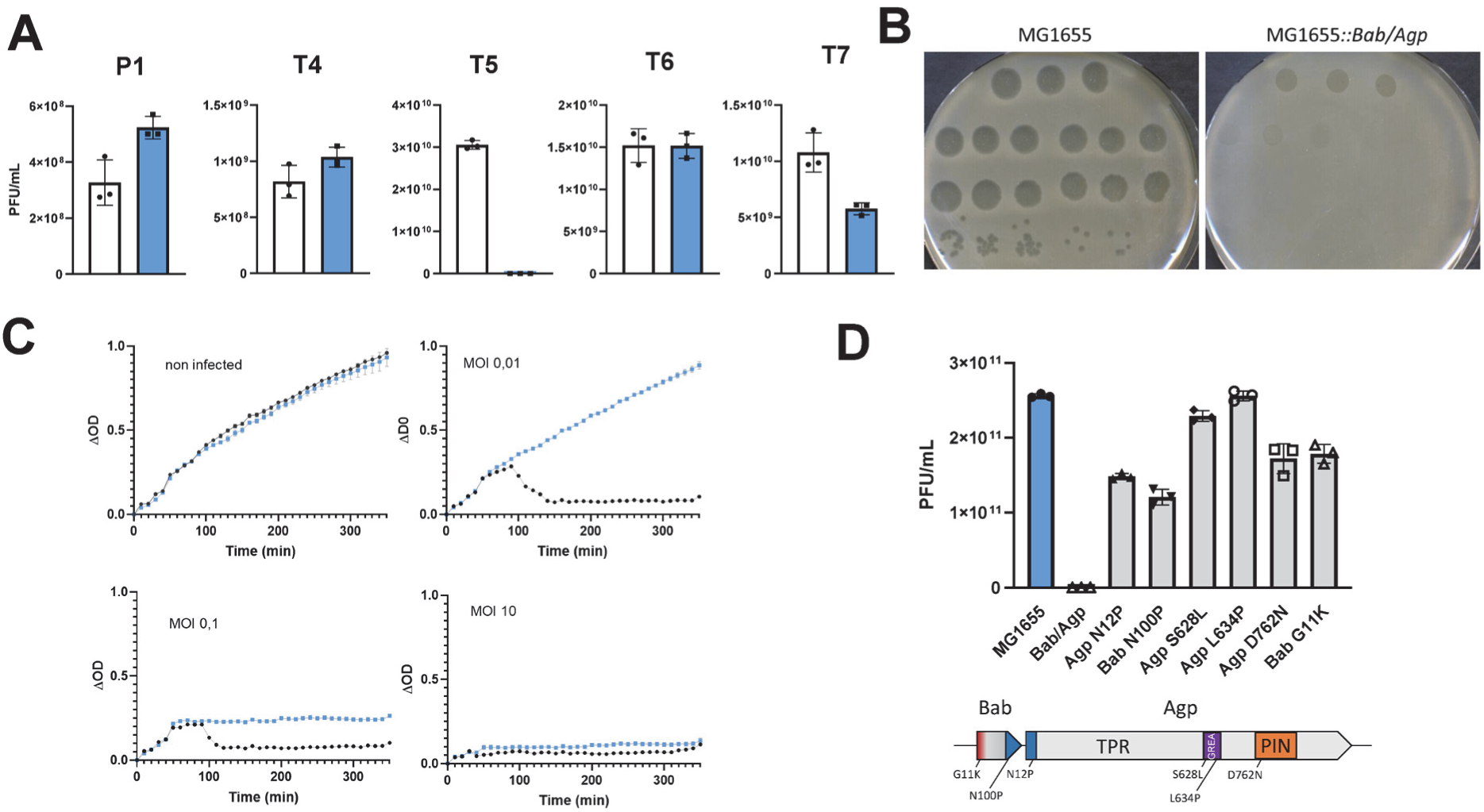
Bab/Agp is an antiphage system. **A.** Plaque counts in MG1655 (white bars) and MG1655*::Bab/Agp* cells (blue bars) infected with different bacteriophages. **B.** Spot assays with 10 fold serial dilutions of T5 suspensions on MG1655 and MG1655*::Bab/Agp* cell lawns. Each dilution was spotted in triplicates. **C.** Growth curves of MG1655 (black) and MG1655::*Bab/Agp* cells (blue) infected with T5 at the given MOI. **D.** Plaque counts in MG1655 and wild-type and mutant MG1655*::Bab/Agp* cells infected with T5. A domain diagram with the position of the tested mutants is given.

To explore whether other T5-like phages are restricted by the Bab/Agp system, we tested seven phages from the subfamily *Markadamsvirinae* (extended data Fig.7A) ^38^. When spotted on the strain carrying the Bab/Agp system, titer reduction was observed for two phages, while two others formed smaller plaques. Three phages appear unaffected, suggesting that not all T5 phages are susceptible to the presence of Bab/Agp. To further explore the specificity of this system, we screened a larger collection of 161 additional phages. For four of them (T145_P1, T145_P3, T145_P4 and T145_P5), no plaques were observed for *Bab/Agp* but the corresponding phages showed only low infectious titer on the MG1655 control strain (extended data Fig.7). The low susceptibility could be attributed to the *Eco*KI restriction system present both in MG1655 and *Bab/Agp*. We deleted the *hsd*R gene of the *Eco*KI system in the *Bab/Agp* strain and retested these four phages (extended data Fig.7B). Strains lacking *hsd*R were now highly susceptible to infection by those phages while strains lacking *hsd*R and containing Bab/Agp were totally resistant in this assay. While belonging to the *Caudoviricetes* class as T5, these bacteriophages belong to the *Drexlerviridae* family instead of the *Demerecviridae* family and have a much smaller genome size (45 kb as compared to 121 kb for T5) thus indicating that the activity of the Bab/Agp system extends across distantly related viruses.

If Abi function of Bab is activated by amyloid signaling, it is expected that β-breaker mutations in the BASS11 motif affect resistance to T5. The β-breaker Bab N100P mutation abolished resistance to T5 (Fig.2D, extended data Fig. 8A). The same was true for a β-breaker mutation at the homologous position in Agp (N12P). A mutation in the predicted N-terminal helix of Bab (G11K) also abolished resistance, indicating involvement of this region in the Abi response. Mutations located in the GreA/B-C domain (Agp S628L, L634P) mimicking mutations altering the GreA/RNA polymerase interaction (extended data Fig.1C) ^39^, both eliminated resistance to phage T5 indicating that integrity of the GreA/B-C domain is required for antiphage activity. A mutation in the predicted catalytic site of the Agp PIN nuclease domain (Agp D762N) ^40^, also led to loss of resistance. For this mutant however, there was a marked decrease in the amount of Agp protein, suggesting that this mutation of the PIN domain compromised global stability of Agp (extended data Fig.8B).

### T5 infection induces Agp-dependent assembly of the BASS11 amyloid motif

The effect of β-breaker mutations in the Agp and Bab BASS11 motifs on antiphage activity suggests that response to T5 involves amyloid conversion of BASS11. We used the BASS11^Bab^-GFP reporter protein to analyze BASS11 assembly following T5 infection. In freshly transformed MG1655*::Bab/Agp* cells expressing BASS11^Bab^-GFP from a plasmid, fluorescence was diffuse in non-infected cells. But 25 min after T5 infection, the fusion protein was assembled into rod-like structures in the vast majority of cells (Fig. 3, extended data Fig. 9). Rod formation did not occur when BASS11^Bab^ N100P-GFP was used as reporter. In size exclusion chromatography, BASS11-GFP was shifted to high-molecular weight fractions upon infection with T5 (extended data Fig.8D). Rod formation occurred independently of full length Bab but was dependent on chromosomally inserted Agp. In particular, assembly was abolished in *Bab/Agp* N12P indicating that the BASS11 motif of Agp is required for rod formation. The S628L mutation in the GreA/B-C domain of Agp also abolished T5-induced BASS11 assembly. These results suggest that during T5 infection an amyloid form of the BASS11^Agp^ can interact with BASS11^Bab^ and convert it to the amyloid fold. Consistent with this model is the observation that synthetic BASS11^Agp^ peptide in amyloid conformation induces insolubility of Bab (but not Bab N100P) when added to crude *E. coli* extracts (extended data Fig.8E).

**Figure 3.**
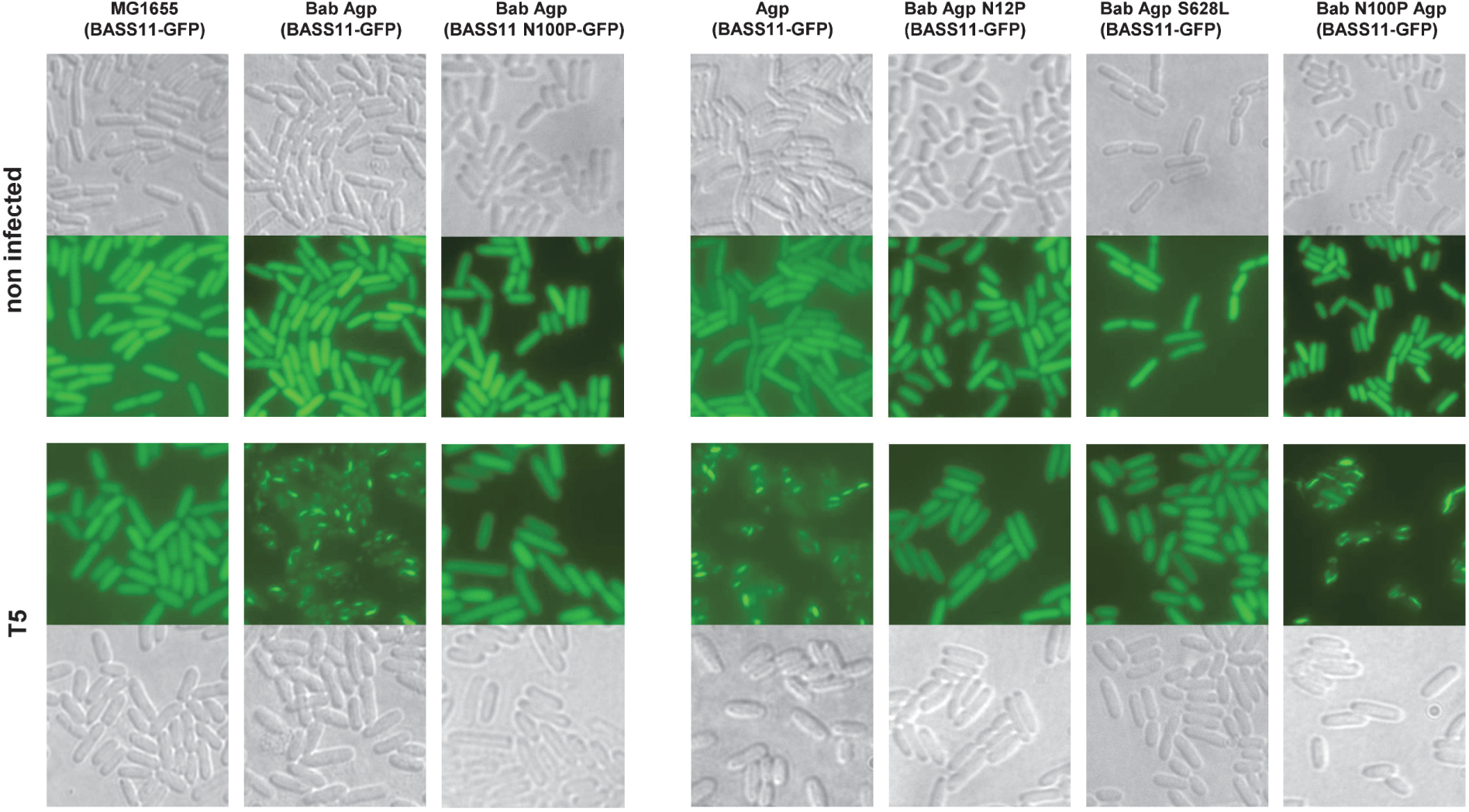
Agp-dependent BASS1-GFP assembly after T5 infection. Cells of the given genotype and expressing either BASS11^Bab^-GFP or BASS11^Bab^ N100P-GFP were infected with T5 at MOI 10 and imaged 25 min after infection. Each panel is 20 x 20 µm.

### T5 infection in *Bab/Agp* cells leads to relocalization of Bab-GFP to the cell periphery and to membrane alterations

We used a Bab-GFP fusion protein as a reporter to analyze localization of full-length Bab upon phage infection. Addition of the GFP tag affected antiphage activity (extended data Fig.8C) but in spite of this limitation we reasoned that this fusion protein might still represent an informative reporter of the behavior of Bab. MG1655*::Bab/Agp* strains were transformed with a plasmid expressing Bab-GFP from the native Bab/Agp promoter. In uninfected cells, fluorescence of Bab-GFP was diffuse. Upon infection with T5, discrete puncta of Bab-GFP (but not Bab N100P-GFP) formed at the cell periphery (Fig. 4, extended data Fig. 10A). Puncta were not formed in a *Bab*/*Agp* N12P background (extended data Fig. 10B). Puncta formation was reduced drastically in *Bab* N100P/*Agp* and *Agp* strains, suggesting that untagged endogenous Bab participates in T5-induced relocalization of the Bab-GFP reporter protein. With the limitation, that the Bab-GFP reporter is not functional in antiphage defense, these observations suggest that a fraction of the Bab protein relocalizes to the membrane region upon infection with T5.

**Figure 4.**
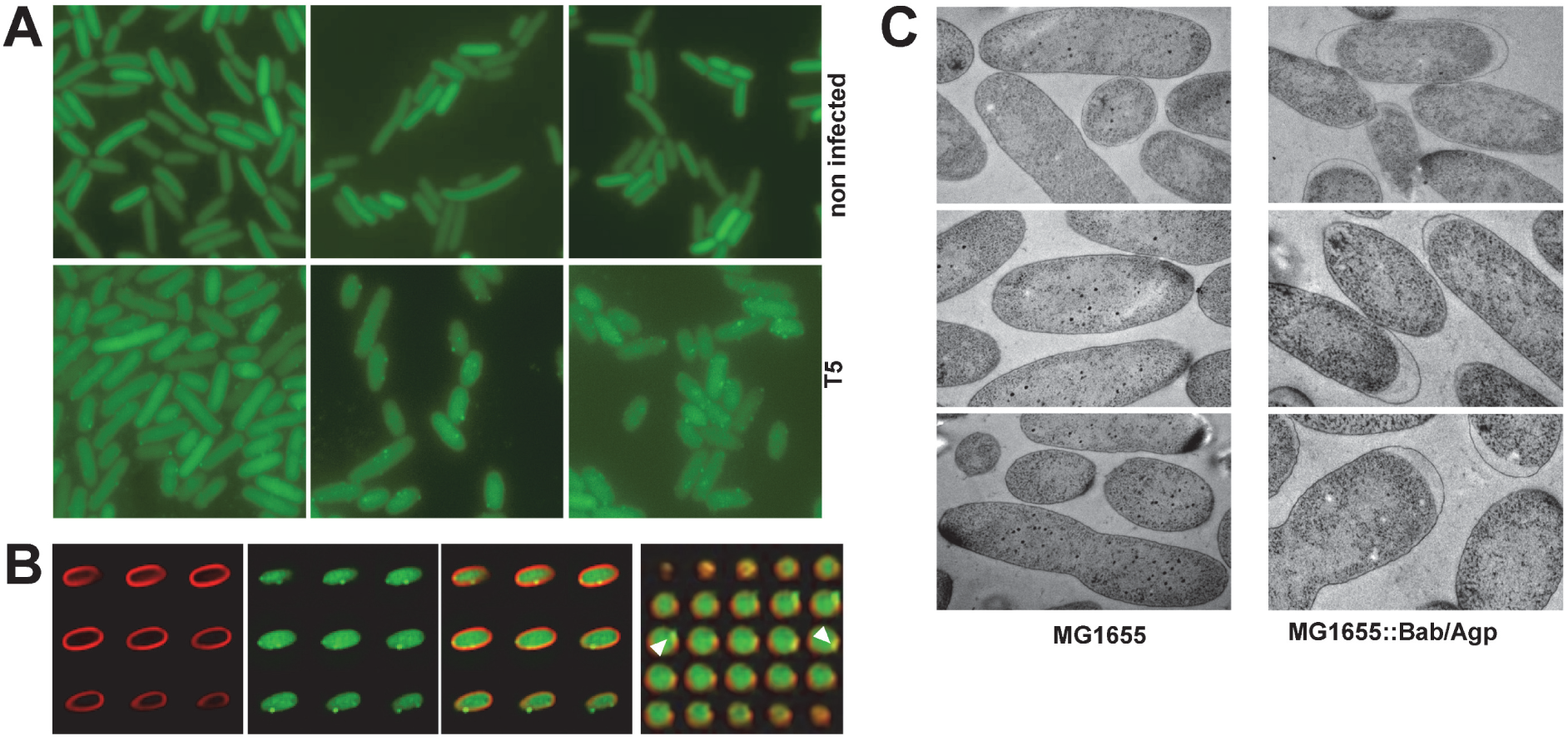
Bab/Agp-dependent Bab-GFP relocalisation and Bab/Agp-dependent membrane alteration during T5 infection. **A.** Bab/Agp cells expressing Bab-GFP where infected with T5 and imaged 25 min after infection. Results with mutant forms of Bab and Agp are given in extended data Fig.10. Each panel is 20 x 20 µm. **B.** Bab/Agp cells expressing Bab-GFP where infected with T5, labelled with Nile Red for membrane staining and imaged 25 min after infection. Panels from left to right are series of z-stacks for Nile red, GFP and overlay. On the right, an orthogonal view of the Nile Red/GFP overlay is given and orthogonal views progress from left to right. The arrowhead point to example of Bab-GFP puncta near the cell periphery. Further images are provided in extended data Fig. 10A. **C.** MG1655 and MG1655*::Bab/Agp* cells were infected with T5 and imaged 25 min after infection. Each panel is 2 x 3 µm. Further images including non-infected cells are provided in Extended data Fig. 11.

We also imaged *Bab/Agp* cells upon infection with T5 using electron microscopy. Twenty five minutes after infection, control MG1655 cells showed electron dense cytoplasmic bodies possibly corresponding to maturating viral precursors, cells appearing otherwise structurally intact (Fig. 4C, extended data 11). In infected MG1655*::Bab/Agp* cells, a retraction of the cytoplasm and formation of a gap between the membrane and the cell wall was observed in the majority of cells (52 of 92 imaged cells, 56%). Membrane retraction was not observed or very rare in infected or non-infected MG1655 and non-infected MG1655*::Bab/Agp* cells (in 0/65, 0/98, 1/87 imaged cells respectively). These observations suggest that a membrane alteration occurs in infected *Bab/Agp* cells undergoing Abi. Together these experiments are consistent with the fact that Bab shows similarity to membrane targeting Hell and Bell-domain proteins (extended data Fig.1A).

### Structure of the globular domain of Bab

We purified the globular domain of Bab overexpressed in soluble form in *E. coli*. Crystals were obtained and the structure was solved by X-ray diffraction. The Bab N-terminal domain displays a compact α-helical fold and is shorter (∼80 amino acids) than previously identified Bell domains which are typically 100 amino acids in length ^23,41^. The fold comprises four helices cupping the N-terminal hydrophobic helix 1 (Fig. 5A). We term this domain Babybell. Helix 1 corresponds to the region of the protein showing the highest conservation in Bab homologs from different enterobacteria. The G11 residue whose mutation (G11K) abolishes Abi-activity is located in the center of helix 1. The experimental crystal structure of Babybell as well as AF models of Bell domains associated to different BASS-motifs in actinobacteria and cyanobacteria, comprise four helices cupping helix 1 (predicted as pore-lining helix by MEMSAT-SVM) (extended data Fig. 12). In spite of the conservation of the N-terminal region (extended data Fig.1A), the folds show extensive variations in the overall topological organization. A possible model for Bab-activation, analogous to the activation of cell death inducing domains of plant ZAR1 ^42^ or fungal HET-S ^13^, could a partial unfolding and the liberation of helix 1 from the helical core leading to membrane targeting and pore-formation.

**Figure 5.**
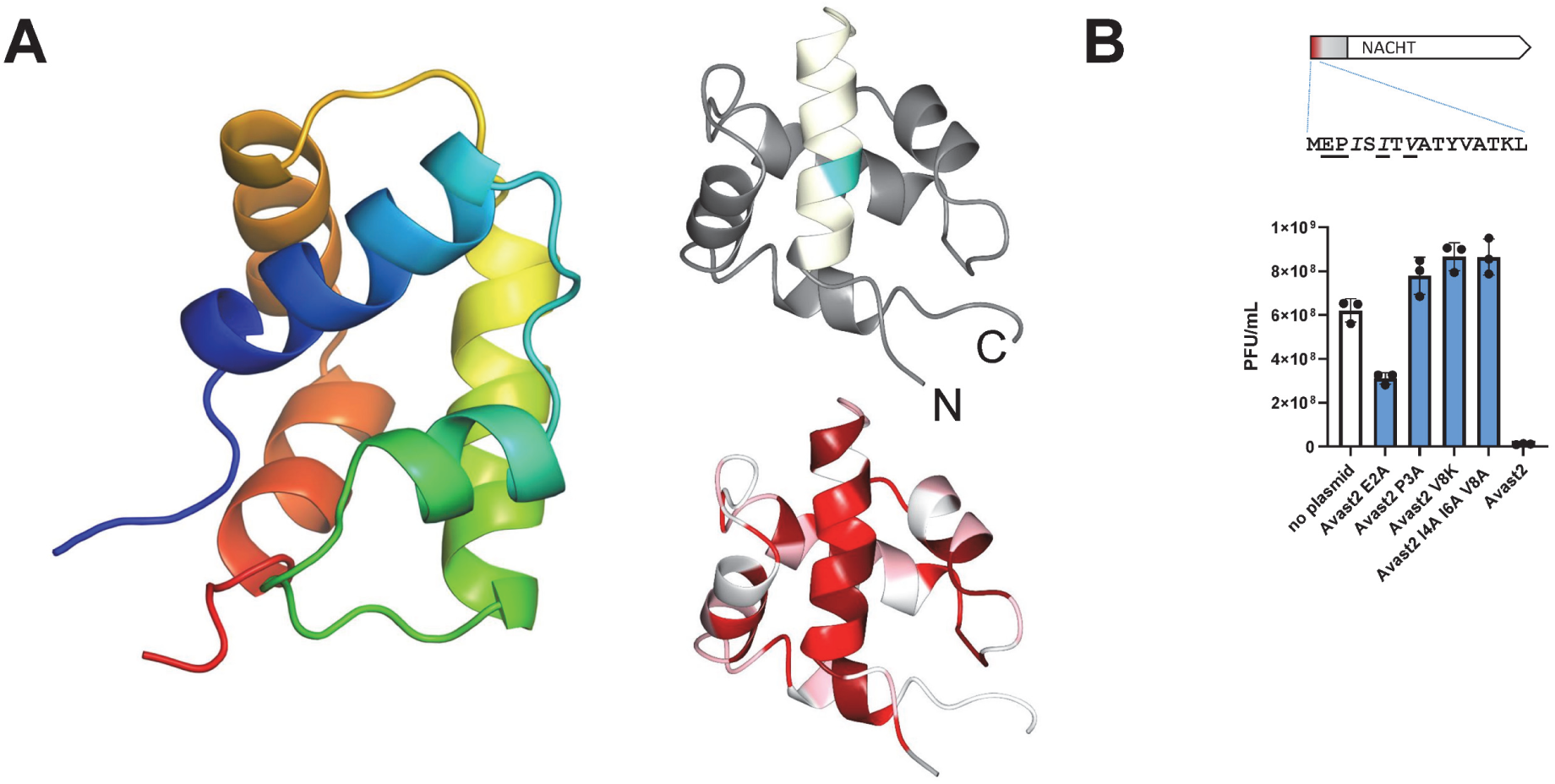
Structure of the Babybell domain of Bab and mutagenesis of the “MEPIS”-motif region of Avast2. **A.** Crystal structure of the globular domain of Bab in rainbow diagram (blue to red from N-terminus to C-terminus). On the left, the top structure is colored with the predicted pore-lining helix in beige and position G11 in cyan. The bottom structure is colored according to sequence conservation in Bab homologs (color code is by quartiles of highest to lowest conservation score, first quartile, dark red, second quartile, red, third quartile dark pink and last quartile, white). **B.** Plaque forming in MG1655 cells expressing wild-type and mutant Avast2 and infected with phage T4. The domain diagram of the Avast2 protein and the position of the mutated sites is given above the graph.

The N-terminal region of Bab common to Hell and Bell domains is also conserved in the N-terminal region of several NLRs with antiphage activity, including Avast2 (extended data Fig.1A), ^28, 29^. In order to start exploring the functional role of this conserved motif in NLR-based antiphage systems, we generated mutations in the N-terminal region of the Avast2 NLR and found that mutations in that region affected its antiphage activity against T4 (Fig. 5B). These results suggest a general conserved role for this “MEPIS”-motif region in cell death induction and antiphage defense.

### Converting a bacterial antiphage defense system into a fungal incompatibility system and vice versa

Our results suggest that Bab functions analogously to the fungal HET-S amyloid-controlled cell death execution protein ^12, 13^. We reasoned that Bab and BASS11 might thus be used to design an artificial fungal incompatibility system akin to HET-S/[Het-s]. As in *E. coli*, a GFP-BASS11^Bab^ fusion protein expressed in *P. anserina* initially displayed a diffuse state (we term [b*]) and then gradually acquired an aggregated foci state ([b]) (extended data Fig.13 and 14A). The [b] state was transmissible to [b*] strains by cytoplasmic contact (extended data Fig.13). N100P and other mutations in conserved key positions of BASS11 (Q92P and D99A) abolished foci formation (extended data Fig.13). When co-expressed, BASS11 motifs of Bab and Agp co-localized in foci (extended data Fig.14B). We conclude that BASS11 can function as amyloid prion forming domain in *P. anserina*. The routine assay for incompatibility is the formation of an abnormal contact line (designated barrage) in the confrontation zone between the tested strains. A barrage formed specifically between Bab and [b] strains while the contact to isogenic [b*] was normal (Fig.6A, extended data Fig. 14C). Microscopic examination confirmed occurrence of cell death in Bab/[b] fusion cells (extended data Fig.14D). [b] prion transmission could be followed using this barrage assay (extended data Fig.13E) ^43^. The Bab/BASS11 interaction also led to sexual incompatibility as described for natural incompatibility systems ^44^ (extended data Fig.13F). Mutations in the N-terminal region of Bab (D2A, G11A/K, Y14A and D15A) and in the BASS11 motif (D99A, N100P) abolished barrage and sexual incompatibility (extended data Fig.13F and G). We conclude that Bab/BASS11 can be tailored into a fungal incompatibility system, analogous to HET-S/[Het-s]. We further reasoned that conversely, it should be possible to design a HET-S-based bacterial Abi system. In the Bab/Agp gene pair, we replaced Bab by *het-S* and the BASS11 motif of Agp by the R0 amyloid motif of NWD2, the NLR receptor paired to HET-S ^19^. *E. coli* strains expressing this chimeric HET-S/R0Agp gene pair on plasmid became resistant to infection by T5 (Fig.6B, extended data Fig. 15A and B). Resistance was slightly less efficient than for Bab/Agp (extended data Fig. 14A). Previously described β-breaker mutations in the HET-S amyloid forming motif (Q240P) or the R0 motif of the R0Agp chimera (H9P) ^13, 19, 22^ abolished resistance (extended data Fig. 14A and B). As in the case of *Bab/Agp* cells (extended data Fig. 5F), cell ghosts were present after infection at high MOI consistent with a mechanism of resistance involving Abi (extended data Fig. 14C). This reciprocal functional repurposing underscores the mechanistic similarities between Bab/Agp and HET-S/[Het-s](NWD2) controlled RCD reactions.

**Figure 6.**
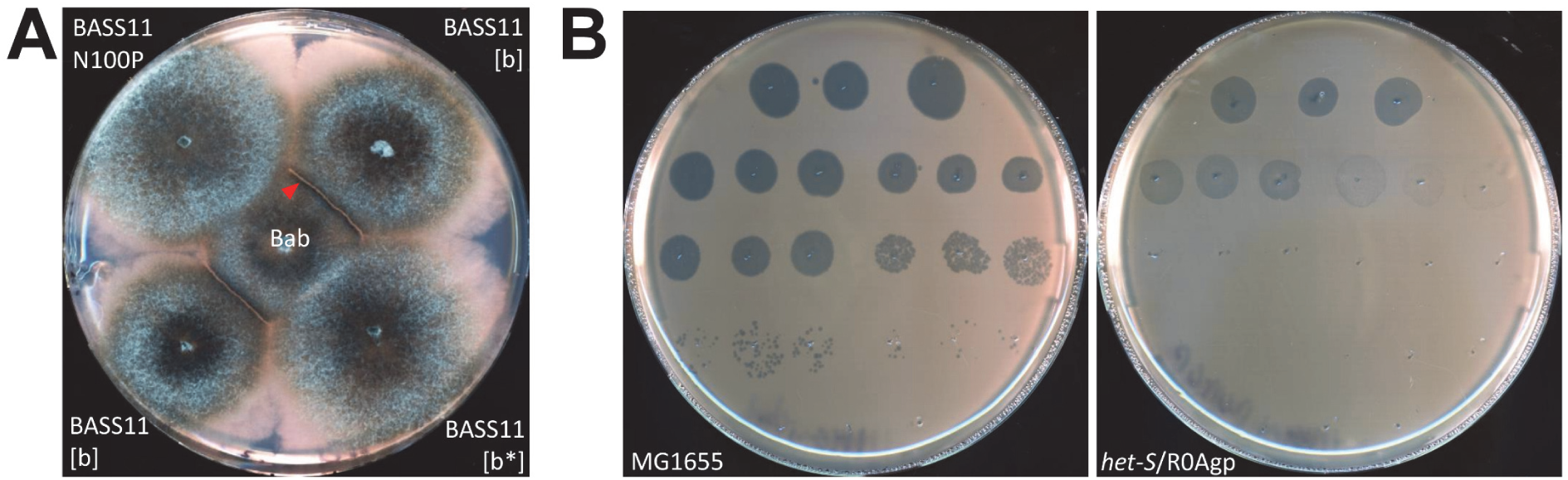
Functional redesigning of Bab and HET-S. **A.** *P. anserina* strain expressing Bab were confronted to strains expressing BASS11 in soluble ([b*]) or aggregated ([b]) state on corn meal agar medium. There is a barrage formation (incompatibility) in the Bab/ BASS11 [b] confrontation, the red arrowhead points to the barrage reaction. **B.** Spot assay with 10-fold serial dilutions of T5 on lawns of MG1655 cells or MG1655 cells expressing *het-S*/R0Agp (see extended data Fig.15 for additional control strains). Each dilution is spotted in triplicate, serial dilution are from top to bottom and left to right.

## Discussion

Building on the work of Kaur and colleagues, our results identify a novel antiphage defense system and lead to a mechanistic model consistent with the previously proposed coupling of TGP proteins to Abi modules ^27^. Bab/Agp confers resistance to T5 and other phages by inducing an Abi response. Abi is dependent on the BASS11 amyloid motif common to Bab and Agp and T5-infection leads to assembly of BASS11. In our model, Abi response is controlled by Agp and executed by Bab, which presumably acts as a membrane targeting protein. As originally proposed, Agp could act as sensor of phage infection by detecting diversion of RNA polymerase activity by phage effector proteins ^27^. Consistent with this model, is the fact that mutations in the GreA/B-C domain affect antiphage activity. Our results, including the functional refurbishing of Bab and HET-S indicate that Agp triggers Bab activation by amyloid templating. In fungal NLR-dependent amyloid signaling, the initial formation of the amyloid fold is controlled by the ligand-induced oligomerisation of the NLR ^19, 20^. Phage infection could lead to Agp oligomerisation and subsequent emergence of the cooperative amyloid fold. Future efforts towards the mechanistic dissection of this Abi system will have to focus on the mechanism of phage sensing and its possible effect on the oligomerisation state of Agp. TGP proteins were proposed to have a dual function, as first line defense systems coupled to back-up Abi modules ^27^. We found no evidence for a Bab-independent activity of Agp against T5 (and T_145 series phages), so a possible function for Agp as first line defense system remains to be established. In the Kaur et al. model, first line defense and Abi are mutually exclusive with Abi activated only in the event of inefficient first line defense. An autonomous Agp activity might only be detectable against phages that do not induce Abi.

While a previous study revealed the existence of NLR/Bell gene pairs in prokaryotes ^23^, the present results now establish a biological role for amyloid signaling in bacteria. Amyloid-controlled immune RCD thus shows conservation in different domains of life from bacteria to fungi and animals. Amyloid signaling adds to the growing list of eukaryotic immune mechanism that are shared with bacteria ^1, 45^. Protease-activated gasdermines are another specific case of RCD system encoded by linked gene pairs and common to bacteria and fungi ^46,47^. Our study expands the functional roles associated to amyloid polymers to antiphage defense ^7^. Amyloid signal transduction is ensured here by a minute sequence motif of only 20 amino acids shared between receptor and effector. This signaling pathway serves as a minimalistic example of immune supramolecular organizing center (SMOC) formation ^48^. We also establish that the Bell domain functions in Abi antiphage defense. This result makes the superfamily of domains including fungal HeLo and Hell, plant CC^HeLo^, animal 4HB of MLKL and bacteria Bell domains an almost universally conserved α-helical membrane targeting module ^14, 23, 49, 50^. Conservation of the “MEPIS” motif in the N-terminal domain of several antiphage NLRs suggests a wider role in antiphage defense ^28, 29^. Sequence and structural homology between these different subfamilies is in some cases exceedingly low and nearly limited to the N-terminal region encompassing the putative pore-lining motif. If the function of the peripheral helices in these domains is merely to act as a sheath cupping the N-terminal helix, this structural task might be ensured by a variety of different sequences. Consequently, prokaryotic and eukaryotic subfamilies could have, over extended evolutionary time, diverged nearly beyond recognition.

The artificial refurbishing of Bab into an incompatibility protein and of HET-S into an antiphage system serves to underline the mechanistic conservation of amyloid RCD signaling in different domains of life. Like mammalian MLKL, both proteins function in amyloid-controlled antiviral RCD. Yet the antiviral strategy differs for fungal HET-S/[Het-s] and bacterial Bab/Agp, the latter system directly kills virus-infected cells while the former is a prophylactic system limiting cytoplasmic exchanges and thus viral transmission between genetically unlike strains.

## Material and Methods

### Sequence analyzes and annotations

Bab homologs were recovered using Jackhhmer at http://hmmer.org/ with default settings, restricting search to prokaryotes and using psi-blast at https://blast.ncbi.nlm.nih.gov with default settings and restricting search to prokaryotes. Sequence redundancy was reduced using DIAMOND-DeepClust and clustering was performed with CLANS in the MPI Bioinformatics Toolkit (https://toolkit.tuebingen.mpg.de/). Sequence alignments were obtained with Clustal Omega at https://www.ebi.ac.uk and visualized with JalView (https://www.jalview.org/) or ESPript 3.0 (https://espript.ibcp.fr). Phylogenetic analyzes were carried out at phylogeny.fr (https://www.phylogeny.fr/). Transmembrane helix predictions were done with MEMSAT-SVM (http://bioinf.cs.ucl.ac.uk/psipred/). Consensus sequence logos were obtained with Weblogo3 (https://weblogo.threeplusone.com). AlphaFold 3 predictions were performed using the AlphaFold server. Structural alignment were obtained at the DALI server (http://ekhidna2.biocenter.helsinki.fi/). Protein structure visualization was done using CCP4MG. Amyloid formation propensity were predicted with ArchCandy 2.0 ^51^ and amyloid co-aggregation using Amylocomp ^52^.

### Strains and DNA constructs

*E. coli* strains used in this study were ECOR25 (provided by the Debarbieux lab), MG1655 (provided by Fabien Darfeuille) and XTL632 (kindly provided by the Court lab). P1, T4, T5, T6 and T7 phage strains were provided by Anaïs LeRhun. The *Podospora anserina* strain used in this study is the previously described *Δhellp (ΔPa_5_8070) Δhet-s (ΔPa_3_620) Δhellf (ΔPa_3_9900)* strain lacking three endogenous proteins with amyloid prion forming domains ^21^. Chromosomal insertion of Agp/Bab derivatives were performed as described ^37^. Chromosomal insertions were verified by PCR and resequenced. A list of all chromosomal insertions in given Table S3. For all constructs, the XTL32 strain was transformed with a PCR fragment generated by an amplification with oligonucleotides 142-149; the resulting product contains a 220 bp and 148 bp region of homology to genomic target insertion site respectively upstream and downstream of Bab/Agp and 336 bp of the Bab/Agp promotor region and 258 bp of the Bab/Agp terminator region. Primers are listed in extended data Table S4. A list of all plasmid construct used in this study is given Table S5. Gene inactivation of the *hsdR* gene with a Kan^R^ cassette was performed as described ^53^.

### Microscopy

Live microscopy was performed with a Leica DMRXA microscope equipped with a Micromax charge-coupled device (CCD) (Princeton Instruments) controlled by MetaMorph 5.06 software (Roper Scientific). The microscope was fitted with a Leica PL APO 100× immersion lens. Fungal cultures were observed directly on solid culture medium and *E. coli* cells were either observed directly on solid growth medium or alternatively liquid cultures were plated on solid medium just before observation. For methylene blue staining, an aqueous 0.5% (w/v) solution was added directly on the mycelium for 1 min, followed by washing with distilled water before observation. *E. coli* cells in the Nile red colocalisation experiment and time lapse microscopy experiments, were observed on a fully-automated Zeiss 200M inverted microscope (Carl Zeiss, Thornwood, NY, USA) equipped with an MS-2000 stage (Applied Scientific Instrumentation, Eugene, OR, USA), a Lambda LS 300 Watt xenon light source (Sutter, Novato, CA, USA), a 100X 1.4NA Plan-Apochromat objective, and a 5 position filter turret. For Nil Red detection we used a Cy3 filter (Ex: HQ535/50 nm – Em: HQ610/75 nm – BS: Q565 nm lp). For GFP displaying, we used a FITC filter (excitation, HQ487/25 nm; emission, HQ535/40 nm; beam splitter, Q505 nm lp). All the filters are from Chroma Technology Corp. Images were acquired using a CoolSnap HQ camera (Roper Scientific, Tucson, AZ, USA). The microscope, camera, and shutters (Uniblitz, Rochester, NY, USA) were controlled by SlideBook software 5.0. (Intelligent Imaging Innovations, Denver, CO, USA). Z-stacks were deconvoluted using the Deconvolution Lab2 plugin. For time lapse microscopy, cells were spread onto a LB medium 2 % agarose pad and imaged every 30min. For membrane staining, cells were incubated before imaging during 30 min with Nile Red (1µg/mL). Negative staining of fibrils was performed as follows. Aggregated proteins were adsorbed onto Formvar-coated copper grids (400 mesh) and allowed to dry for 15 min in air; grids were then negatively stained for 1 min with 10 μl of freshly prepared 2% uranyl acetate in water, dried with filter paper, and examined with a Hitachi H7650 transmission electron microscope (Hitachi, Krefeld, Germany) at an accelerating voltage of 120 kV at the Pôle Imagerie Électronique of the Bordeaux Imaging Center using a Gatan USC1000 2k × 2k camera. Electron microscopy of *E. coli* cells was performed as described ^54^.

### ThT and Congo Red binding assays

Thioflavin T (Sigma) was used at 50 µM with 80 µM of synthetic peptides in a volume 100 µl in 20 mM Tris HCl pH 8. Fluorescence was measured in a Clariostar plate reader with excitation at 440 nm and emission at 490 nm. Congo Red (Sigma) was used at 15 µM with 30 µM of peptide in 20 mM Tris HCl pH 8. Absorbance spectra between 400 and 600 nm were recorded using a Nanodrop 1000 spectrophotometer (Thermo Scientific).

### Phage assays

For plaque formation assays, 100 uL of phage suspensions at 10 pfu/mL are added to 1 ml of overnight cultures of *E. coli* strains in 2xYT at 37°C with agitation and 4ml of preheated top agar are added (LB with 0.1mM MnCl_2_, 5 mM MgCl_2_, 5 ml CaCl_2_, 7.5 g/L of agar) and the mixture is poured on LB solid medium (with appropriated antibiotics when plasmid containing strains are used). Plates are incubated from 6 to 16h at 37°C. For direct spot tests, plates with cell lawns in top agar are obtained as above and 4 µL of 5 serial 10-fold dilutions of phage suspensions starting with 10^2^ dilutions of high titer phage stocks at ∼10^8^ pfu/mL. (in LB with 0.1mM MnCl_2_, 5 mM MgCl_2_, 5 ml CaCl_2_) are spotted in triplicates. Plates are incubated from 6 to 16h at 37°C. For phage assays in liquid cultures, exponentially growing cultures are diluted in LB with 0.1mM MnCl_2_, 5 mM MgCl_2_, 5 ml CaCl_2_ to an OD of 0.1 and 20 µL of phage suspensions are added to 200 µL of bacterial culture inoculated in a microwell plate and incubated at 37°C in a Bioscreener SW (ThermoScientific). For the *het-S*/R0Agp construct, phage assays after growing cells for 7 hours after plasmid transformation until OD reached 1. The screen of 161 phages from Debarbieux lab was performed by spotting a single drop (4 µL) of each phage stock on a LB plate covered with a lawn of each bacterial strain seeded from an exponentially growing culture in LB and plates were incubated overnight at 37°C. Clear spots indicated which phage that was subsequently tested to determine the EOP (efficiency of plaquing) using the same process but plating 4µL drops of 10-fold serial dilutions. Phages T145_P1/3/4/5 were isolated from sewage using strain T145 and their respective accession numbers are OZ035767.1, OZ035793, OZ035765.1 and OZ035786.1 .

### Fungal assays

DNA transformations were performed as previously described ^55^ using pBC1004 bearing the *hph* gene conferring resistance to hygromycin in co-transformation ^56^. All fungal cultures were performed on solid medium expect for protein extraction. Barrage assays were performed on solid corn meal agar medium (D0) and synthetic medium (SU), crosses were performed on synthetic M2 medium ^57, 58^. Full length Bab and the BASS11 motif of Agp and Bab were cloned as GFP or RFP fusion proteins under dependence of the strong constitutive GPD promotor as previously described ^21^. For foci formation assays, a collection of primary transformants expressing GFP-Bab(86-109) or GFP-Bab(86-109)mutants were analyzed by fluorescence microscopy 5,7,11 and 30 days after transformation for foci formation. For foci transmission assays, primary transformants were confronted with a donor strain expressing GFP-Bab(86-109) foci and were further subcultured 2 days after contact with the donor strain analyzed by fluorescence microscopy for foci formation. For fertility assays, strains were crossed by confrontation on M2 medium and number of perithecia were counted under a binocular lens after 7 days.

### Western-blotting and antibodies

For *Podospora* crude extracts preparation, for each strain, a dozen implants of mycelium were grown in 5 ml 2xYT medium containing 10 μg/ml ampicillin at 26°C for 2 days with agitation. After pouring out the medium, mycelial mats were washed in 5 ml distillated water and excess liquid was removed on Whatman 3MM paper and the mycelium transferred in beads-containing tubes (Lysing Matrix A, 6910100, MP Biomedicals) with 300 μl lysis buffer (50 mM Tris pH 8, 150 mM NaCl, 1 % SDS, 100 mM DTT, 10 M urea). Cells were lysed in a FastPrep instrument (MP Biomedicals) for 2 rounds of 30 seconds at 6 m/s. Samples were centrifuged at 17 000 g for 5 minutes and supernatants were collected. Samples were analyzed in 6 % Tris-glycine SDS-polyacrylamide gels, transferred on nitrocellulose membranes (Protran 0.45 μm, Amersham) using a semi-dry cell (Trans-Blot SD, Bio-Rad) in transfer buffer (39 mM glycine, 48 mM Tris pH 8.3) at 10 V for 30 minutes. Membranes were blocked in TBS-T buffer (20 mM Tris pH 7.6, 137 mM NaCl, 0.1 % Tween 20) containing 5 % (w/v) nonfat dried milk for 1 hour, incubated in a 1:1000 dilution of the primary antibody (either anti-V5 tag mouse monoclonal antibody, MCA1360, Bio-Rad, or anti-Softag1 SLAELLNAGLGGS mouse monoclonal antibody to detect the β’ subunit of *E. coli* RNA polymerase, NT73, AntibodySystem, or anti-GFP mouse monoclonal antibody, 11814460001, Roche) for 1 hour. After 30 minutes of washing in TBS-T buffer, membranes were incubated in a 1:10 000 dilution of the secondary antibody in TBS-T buffer (sheep anti-mouse peroxidase antibody, A5906, Sigma-Aldrich). After washing, membranes were probed with a chemiluminescent substrate (Clarity Max Western ECL Substrate, Bio-Rad) and analyzed in a ChemiDoc imaging system (Bio-Rad). SEC fractionation of *E. coli* crude extracts was performed on a Superdex 200 increase 10/300 GL using 500 µL of crude extracts and 1 ml fractions

### Protein purification

Bab(1-90) was cloned into a pET24a+ expression plasmid between *Nde*I and *Xho*I restriction sites. 100 ng of pET24a+_BABYBELL plasmid was transformed into *Escherichia coli* BL21(DE3)pLysS competent cells. Cells were spread onto an agar-plate containing kanamycin. The next day, a single colony was picked and inoculated into a preculture of LB medium. After overnight incubation at 37°C, 220 rpm, 1 liter of LB culture implemented with kanamycin, was inoculated with the preculture at a rate of 0.1 final OD. The culture was incubated at 37°C, 220 rpm for cell growth. When the exponential phase is reached, protein expression was induced with 1 mM IPTG. After overnight incubation at 28°C, 220 rpm, cells were harvested at 6,000 g, 15 minutes. The bacterial pellet was resuspended in 80 mM NaCl, 20 mM imidazole, 20 mM HEPES, pH 7.5 (lysis buffer) then lysed using a sonicator (Sonoplus US 70/T, Brandelin). Cell debris and insoluble proteins were removed by centrifugation at 20,000 g 30 minutes, 4°C. The supernatant was applied to a Ni-NTA-affinity column (HisTrapHP, GE Healthcare) previously equilibrated with the lysis buffer and the column washed with the same lysis buffer until an OD of 5 mOD is obtained. Proteins were eluted using a linear gradient up to 50 % of 80 mM NaCl, 1 M imidazole, 20 mM HEPES, pH 7.5 (elution buffer) over 15 column volumes (CV) followed by an isocratic step gradient of 100 % elution buffer over 2 CV. Selected fractions were dialyzed 3 times for 1 hour against the same imidazole-free buffer. The His-tag was then cleaved off by TEV protein, added in a 3:100 ratio (w/w) in buffer 80 mM NaCl, 1 mM DTT, 0.5 mM PMSF, 0.5 mM EDTA, 0.02 % (w/v) azide, 20 mM HEPES, pH 7.5 (dialysis buffer), and incubated overnight at room temperature under gentle agitation. The cleaved protein was purified by a second nickel affinity chromatography. This HisTrapHP column was equilibrated with the dialysis buffer. The column was washed with dialysis buffer until an OD of 5 mOD is reached. The elution was performed with a first isocratic step gradient at 50 % of dialysis buffer containing 1 M imidazole on 2CV and a second isocratic step gradient at 100 %. A final SEC step was performed to obtain the pure and monodisperse protein. The unretained fraction of the Ni-NTA column is deposited on a HiLoad 16/600 Superdex 75 (Cytivia) previously equilibrated with 80 mM NaCl, 20 mM HEPES, pH 7.5. The protein was eluted at an elution volume compatible with a molecular weight of the protein in a narrow and symmetrical peak. The protein was concentrated at 60 mg/ml using centrifugal filter devices (3K Amicon Ultra-4, Millipore). Around 12 mg of protein per liter of culture was obtained with a purity of 99 %, as judged by SDS gel electrophoresis with Coomassie staining. The recombinant pET24-Bab(50-109)-6His plasmid was transformed into E. *coli* lemo 21 (DE3) cells for protein expression. After transformation by electro-shock, the cells were recovered in LB medium and plated on 30ug/mL of kanamycin, 30ug/mL chloramphenicol-containing LB agar plates, followed by overnight incubation at 37°C. Colonies were selected and inoculated in 5 mL of LB containing Kanamycin (30 µg/mL) and chloramphenicol (30 µg/mL), grown at 37°C with shaking at 200 RPM for a few hours. These cells were then recovered by centrifugation and inoculated into 100 mL of M9 minimal medium enriched with, D-Glucose labeled with ^13^C6 (2 g/L) and ^15^NH4Cl (1 g/L), MEM vitamins, and essential salts (MgSO₄, ZnCl₂, FeCl₃, CaCl₂), kanamycin (30 µg/mL) and chloramphenicol (30 µg/mL). After overnight growth at 37°C and 220 RPM, the culture was diluted into 1 liter of the same previously described M9 media, cultured until OD600 reached 1 and induced with 1mM IPTG, at 30°C overnight, 220 RPM. Post-expression, cells were harvested by centrifugation (7000 g, 30min) and lysed by sonication on ice in 25 mL buffer A (Tris 50 mM pH 8, 150 mM NaCl). The cell suspension was centrifuged (15 000 g ,1 h, 4°C) to collect inclusion bodies. The pellet was then resuspended in 15 mL of extraction buffer B (50 mM Tris pH 8, 0.5 M NaCl, 6M Gu-HCl). The suspension was incubated overnight at 60°C, sonicated and ultra-centrifuged (250 000 g, 1 h, 25°C) before purification steps. Protein was purified over a 5 mL Histrap HP column (Cytiva), previously equilibrated with buffer B1 (50 mM Tris pH 8, 0.5 M NaCl, 20 mM imidazole, 8 M urea,). Protein was eluted with a step of 70% of buffer B2 (50 mM Tris pH 8, 0.5 M NaCl, 500 mM imidazole, 8 M urea), monitored by UV absorbance at 280 nm, on Akta systems. Protein was further buffer exchanged into 175mM acetic acid pH 2.5, over a HiPrep 26/10 desalting column (Cytiva) and pure fractions were analyzed on 12% Tris-tricine SDS-PAGE. For assembly, protein aggregation was initiated by raising the pH to 8 with 12% (v/v) 3 M Tris-HCl (pH 8) and agitated at room temperature for several days. For purification of full length Bab, the pET-24 Bab 6his plasmid was transformed into BL21(DE3) pLysS cells. An overnight preculture from a single colony was inoculated in 50 ml of 2xYT with kanamycin (50µg/mL) and chloramphenicol (34 ug/mL) and grown under strong agitation at 37°C until OD_600nm_ reached 0.6, ITPG to 1 mM was added and the culture grown for 6 hours at 37°C. Cells were pelleted, frozen at 80°C and suspended in 2 ml of 150 mM NaCl Tris 50 mM pH 8 and sonicated on ice. The crude extract was centrifugated for 30 min at 17 000g and resuspended in 6 M GuHCl 150 mM NaCl Tris 50 mM pH 8 and purified with 600 µL bulk Talon resin (Takara). Proteins were eluted in 1 ml of 200 mM imidazole 8 M urea 150 mM NaCl Tris 50 mM pH 8 and dialyzed against water overnight. Bab 6his assembly occurred during dialysis.

### Crystallization of Bab(1-90), Data collection, structure solution and refinement

Crystallization screening were performed using Jena Science and Nextal screens by sitting-drop vapor-diffusion technique in a 96-well format crystallization plate. The final volume of the drop was 0.4 µl, with 0.2 µl of BABYBELL at 10, 15 or 18 mg/ml and 0.2 µl of the reservoir solution. The crystallization plates were made using the Mosquito robot. The plate was incubated at a constant temperature of 20°C. The crystallization condition D4 of JCSG++ HTS screen (30% (w/v) PEG 8,000, 100 mM sodium acetate pH 4.5, 200 mM lithium sulfate) that gave the best crystals. Protein crystals grew in 5 days and are diamonds with dimensions of 110*325 µm. Protein crystals belong to the monoclinic C2 space group with cell parameters: a=108.04 Å b=119.63 Å c=64.17 Å β=92.5° (see Table S6 for statistics of data collections). The asymmetric unit (ASU) is composed of 8 independent monomer copies. The diffraction data have been collected up to 2.4 Å resolution at 120K on a home source Rigaku FRX rotating anode equipped with High flux confocal mirrors and a pixel Hybrid HyPix6000 detector. Indexing and integration of the diffraction intensities were performed with CrysalisPro (version 1.171)^59^. Molecular replacement was performed using PHASER ^60^ with a search model generated by Alphafold (https://alphafold.ebi.ac.uk). The Structure was completed and improved in COOT ^61^ before refinement with BUSTER ^62^. Models of the 8 molecules in the ASU were optimized through iterative rounds of refinement and model building. Model quality was validated using MolProbity ^63^. Crystal packing was examined using PISA ^64^. Refinement statistics are included in the mmCIF file deposited to the PDB under the code 9H9V.

### Solid-state NMR of Bab(50-109)

Bab (50-109) fibrils were packed in a 3.2 mm rotor. 83 kHz ^1^H decoupling was used during indirect and direct acquisition times. Temperature was calibrated according to the water proton resonance and set to 278 K ^65^. Chemical shifts were calibrated according to DSS. Two-dimensional proton-driven spin-diffusion experiments were recorded at 800 MHz ^1^H frequency at a magic-angle spinning frequency of 11 kHz, using 50 ms and 150 ms mixing times for ∼3 days and ∼4.5 days respectively. An initial ^1^H-^13^C cross-polarization mixing of 1 ms was used. The two-dimensional INEPT experiment was recorded for ∼18 hours. Spectra were analyzed using CCPNMR. Secondary chemical shifts were calculated according to Luca et al. ^31^.

---

**Extended data Figure 1.**
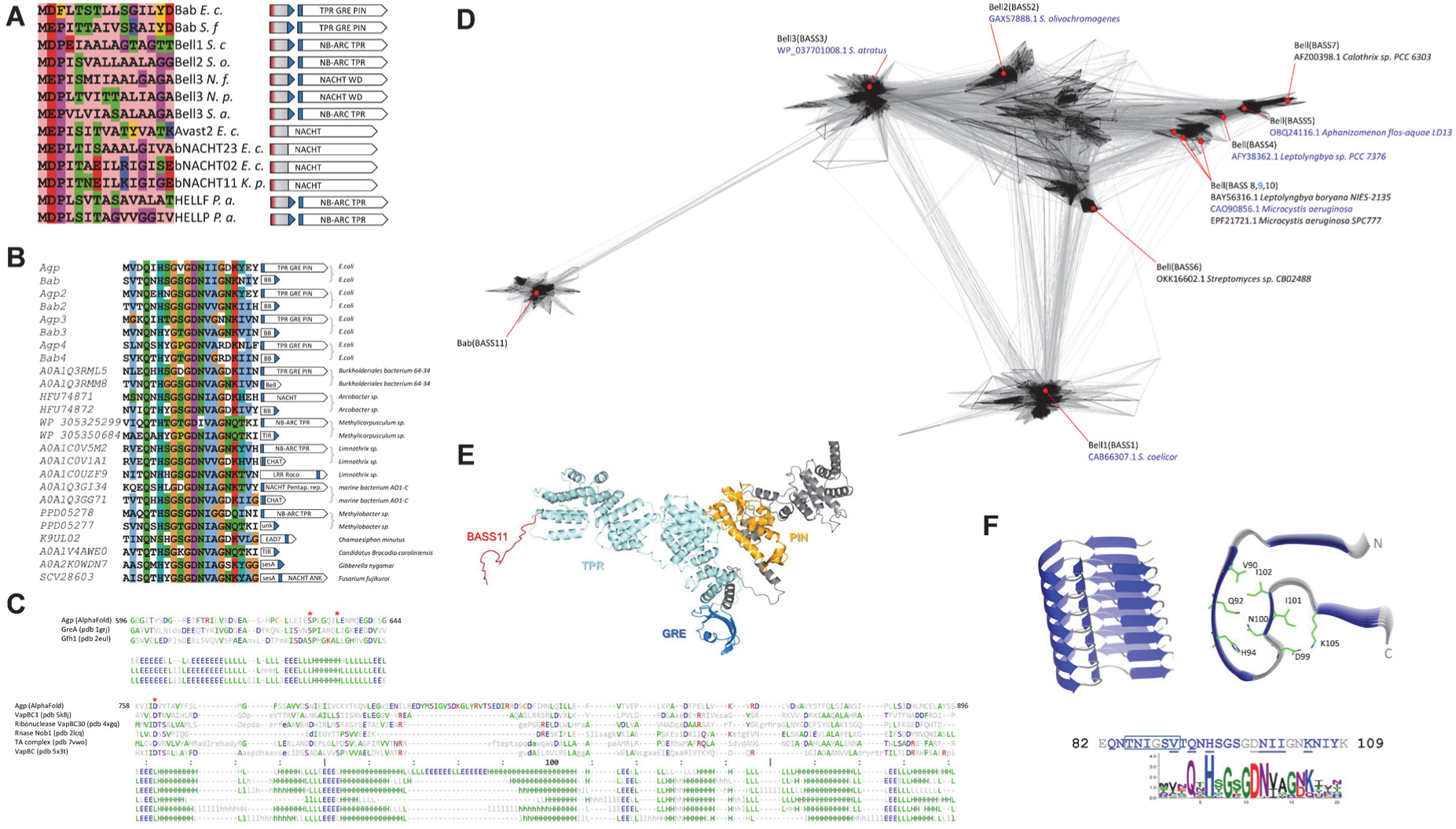
Sequence analysis of Bab/Agp. **A.** An alignment of the sequence of the N-terminal region of selected bacterial Bab and Bell domain proteins, fungal Hell domain proteins and Avast2, bNACHT02, 11 and 23 NLRs involved in antiphage defense is given together with the domain and gene architecture in which the sequences occur. The color code is as in Fig.1A, with amyloid forming sequences in blue and predicted N-terminal hydrophobic helix in red. Genbank entries for all mentioned gene products are given in extended data Table S1. **B.** Domain associations of BASS11-motifs. An alignment of BASS11-motifs occurring in different domain architectures is given. Corresponding domain architectures and the motif position (in blue) are given for each protein. Sequences grouped with a bracket accolade are encoded by adjacent genes. **C.** Upper panel is a structural alignment of the PIN domain region of Agp with selected PIN domain proteins as given by DALI. The red asterisk mark the position of a predicted catalytic site residue. The lower panel is a structural alignment of the GreA/B- C region of Agp with GreA and Gfh1 as given by DALI. The red asterisk mark positions of residues whose mutation affects binding to RNA polymerase. **D.** Clustering of bacterial Bell domain proteins. The N-terminal region of Bab corresponding to the predicted helical domain (1-80) was used in psi-blast and HHMER searches to recover homologous sequences. A total of 2267 non-redundant sequences were clustered with CLANS using default settings. Domain organization and accession numbers are given for selected sequences representing the different classes of BASS-motifs. Sequences do not include the BASS-amyloid motif region so homology in the BASS-region does not contribute to clustering. The sequences given in blue are those for which an AlphaFold model is given in extended data Figure 12. **E.** AlphaFold model of Agp. The TPR, GreAB-C and PIN domain are given in cyan, blue and gold respectively. **F.** AlphaFold 3 model of the Bab(81-109) sequence in multimeric form (with a sequence input of 7 monomers), the positions of predicted β-strands are given in blue in the amino acid sequence. A consensus sequence logo of the BASS11 motif based on the alignment of extended data figure 1B is given. Position of conserved residues underlined in the sequence are shown on the structural model. The boxed region corresponds to residues with spectral assignment in ssNMR for which chemical shift and thus secondary structure have been determined.

**Extended data Figure 2.**
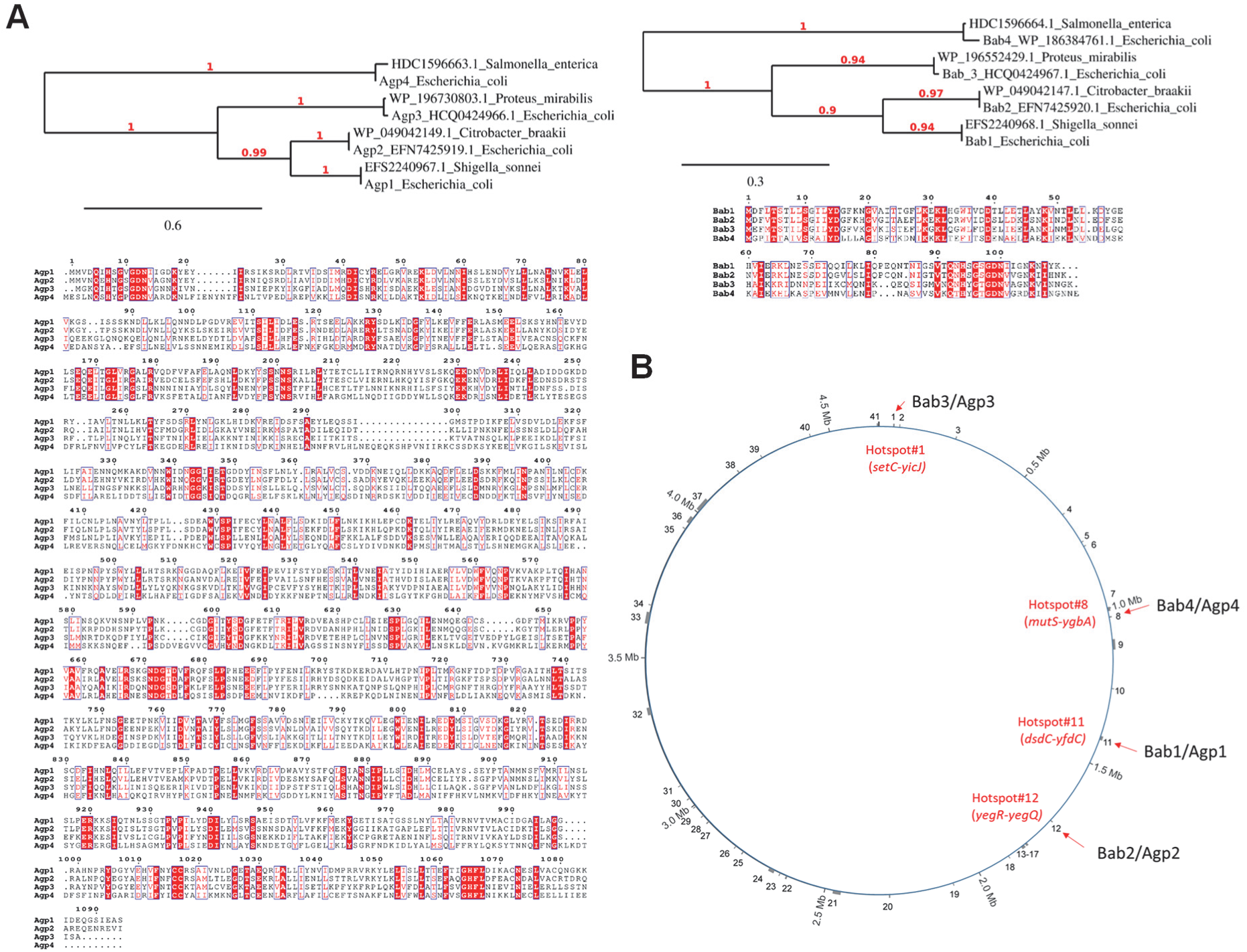
Bab/Agp paralogs and their genomic location. **A.** Phylogenetic tree and sequence alignment of Bab and Agp protein sequences showing four classes of paralogs. Trees were constructed with a representative *E.coli* sequence and an orthologous sequence from a distinct species. Note that the Agp and Bab tree are congruent. **B.** Map of the *E. coli* chromosome with location of 41 defense islands modified from Hochhauser et al (2023), the chromosomal position of the different Bab/Agp paralogs is given.

**Extended data Figure 3.**
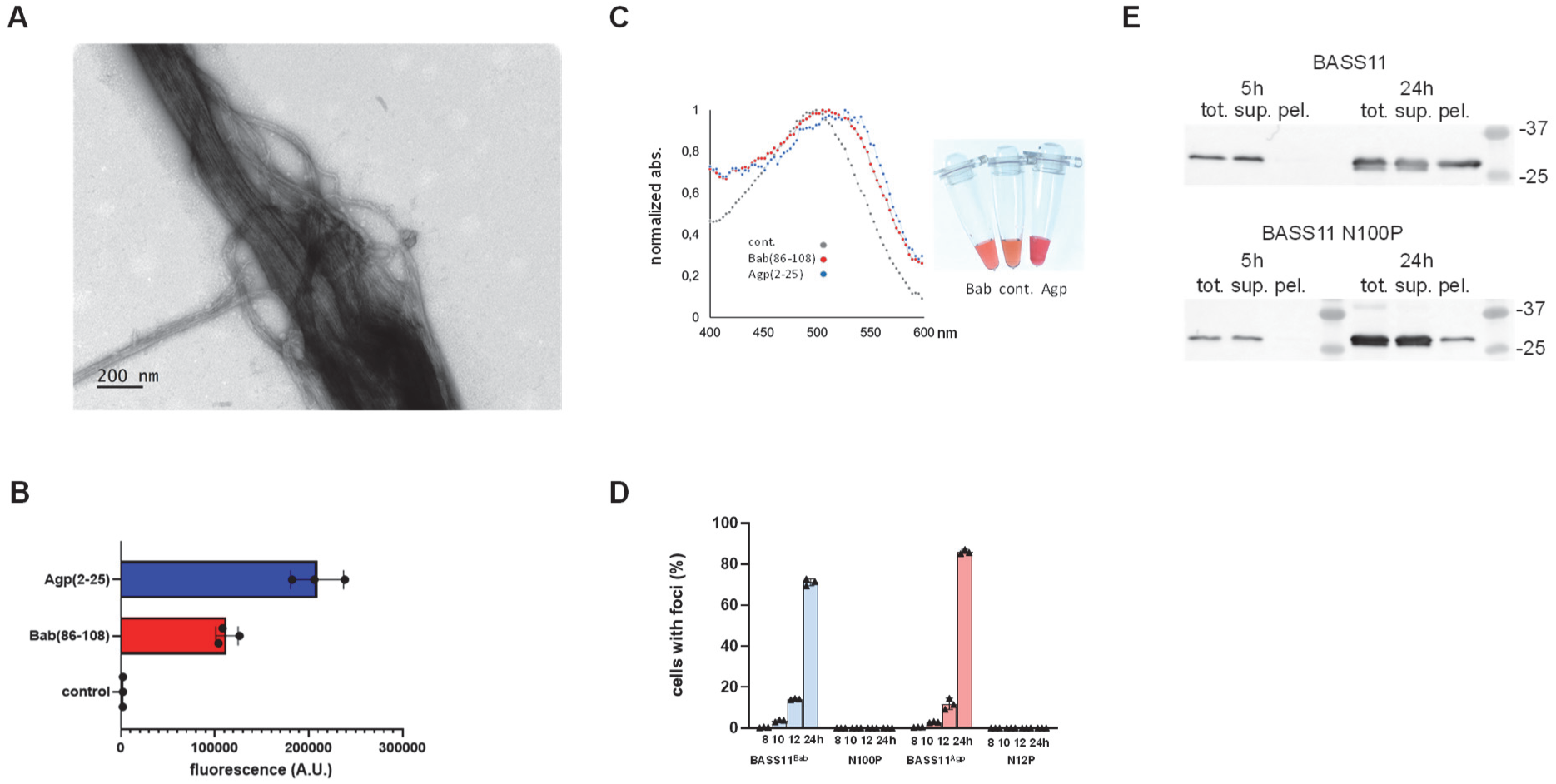
Properties of the BASS11 amyloid motif. **A.** Electron micrographs of full length Bab-6his fibrils. **B.** Thioflavin T fluorescence induced by Agp(2-25) and Bab(86-108) fibrils measured at 50 µM ThT with 80 µM of synthetic peptide. **C.** Bathochromic shift in the Congo red absorption spectrum in the presence of Agp(2-25) or Bab(86-108) fibrils measured at 15 µM Congo red and 30 µM of synthetic peptides, absorbance are normalized with the absorbance value at the maximal absorbance wavelength (respectively 499 nm for free CR, 505-511 nm for CR with Bab(86-108) and 526 nm for CR with Agp(2-25)). **D.** The graph gives the proportion of cells containing GFP foci at different time points after transformation for the listed strains expressing wild-type or mutant versions of the BASS11 motif fused to GFP (BASS11-GFP). **E.** Insolubility of BASS11-GFP in cells with foci. Anti-GFP western-blot of crude and fractionated *E. coli* extracts. Cells were harvested 5h or 24h after transformation and crude extracts (tot.) were centrifuged for 1h at 17,000g to yield the supernatant (sup.) and pellet (pel.) fractions. Size of molecular weight markers is given in kDa.

**Extended data Figure 4.**
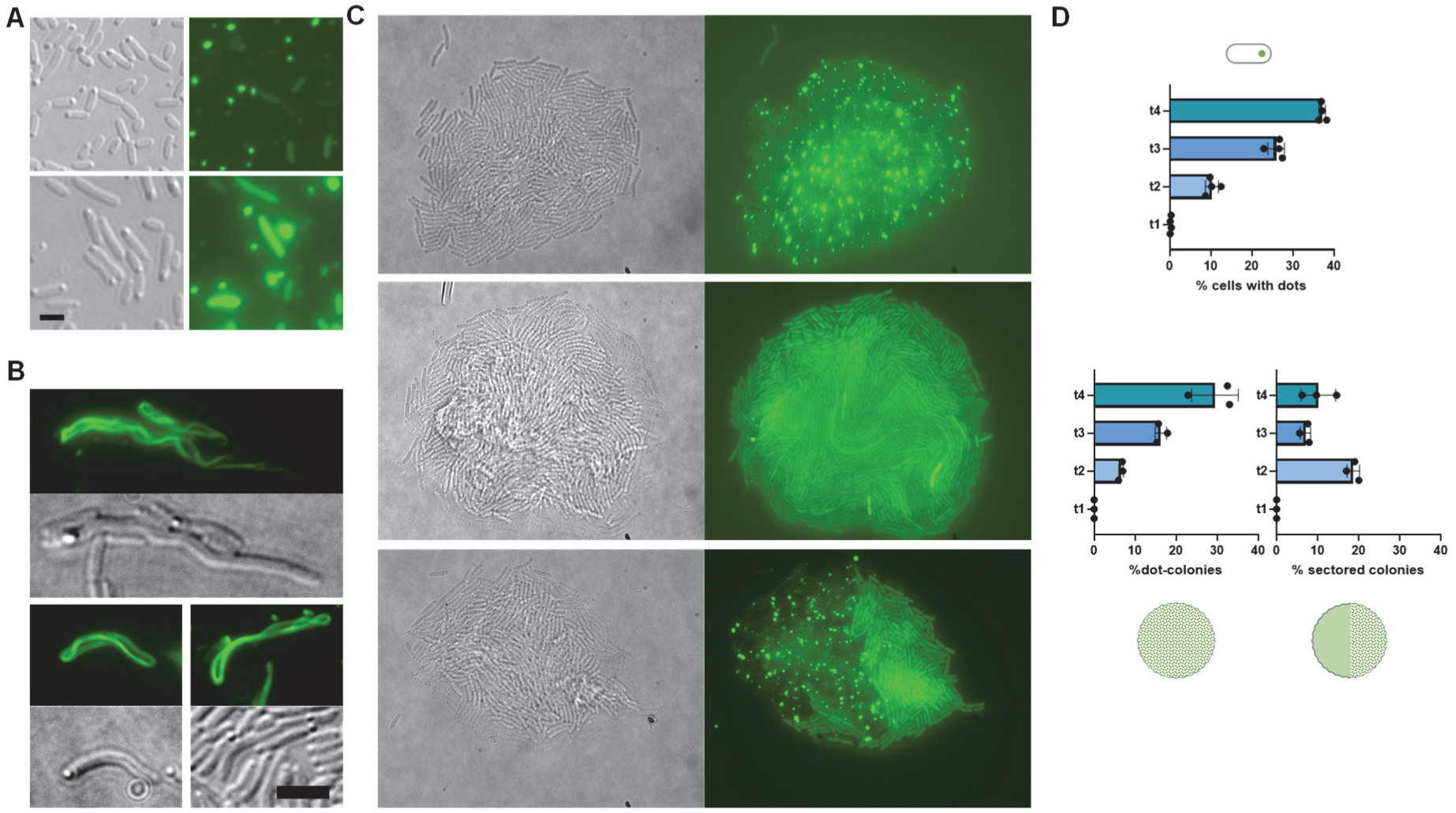
Formation of dot-like foci by BASS11-GFP in *E. coli.* **A.** Colocalization of BASS11-GFP foci with refringent bodies at the cell poles, scale bar is 2 µm. **B.** Examples of cells with BASS11-GFP cable-like structures, scale bar is 4 µm**. C.** Representative examples of colonies obtained after plating liquid cultures containing cells with dots and cells with diffuse BASS11-GFP fluorescence: from top to bottom, a colony homogeneously displaying cells with dots, a colony homogeneously displaying cells with diffuse fluorescence and a chimeric sectored colony with a dot and a diffuse fluorescence sector. Colonies were imaged after 6 hours of growth at 37° on LB medium. Images are 130 x 100 µm in size. **D.** Correlation between the proportions of dot cells in starting liquid cultures and dot colonies formed after plating. The upper graph gives the proportion of cells with dots at four different time points after transformation of MG1655 *E. coli* cells with a plasmid expressing BASS11-GFP. The two lower graph give the proportion of dot and chimeric sectored colony obtained after plating the liquid cultures at time point t1 to t4.

**Extended data Figure 5.**
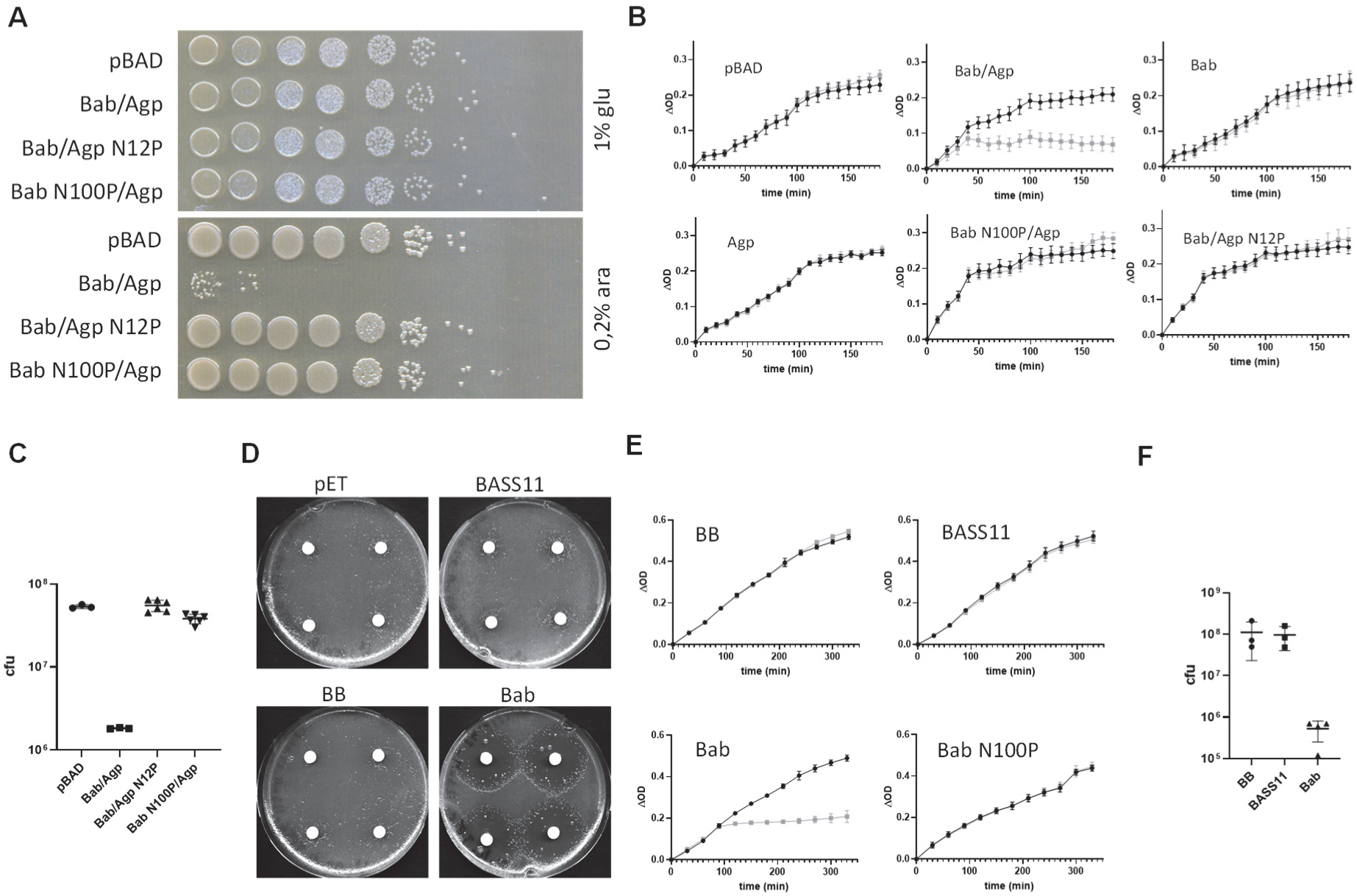
Inducible expression of Bab/Agp is toxic. **A.** Drop test of 10-fold serial dilutions of *E. coli* TOP10 cultures containing a pBAD plasmid with Bab/Agp constructs as given on 2xYT Amp medium with 1% glucose or 0.2% arabinose after 16h incubation at 37°C. **B.** Growth curves in liquid cultures of *E. coli* TOP10 cells containing a pBAD plasmid with Bab/Agp constructs as given, cultures were shifted to 0.2% arabinose medium at time zero and had a starting OD_600nm_ of 0.1. Results are presented as OD variation (ΔOD), the OD value at time zero have been substracted for each culture. The grey curves are in 0.2% arabinose, the black curves in 1% glucose. **C.** Colony formation of *E. coli* TOP10 cells containing a pBAD plasmid with Bab/Agp constructs as given. Cells were shifted to arabinose 0.2% medium for 3 hours and plated on glucose 1% medium. Results are given as cfu per mL of cultures with an OD_600nm_ of 1. **D.** Toxicity of IPTG inducible expression from pET24 vectors in *E. coli* BL21(DE3) pLysS cells containing Bab derived constructs. Cells laws (∼10^6^ cells per plate) were overlaid with a filter paper and 25 µL of IPTG at 125 mM were added on the filter paper. Plates were incubated for 30 hours at 26°C. Bab corresponds to full-length Bab, BASS11 to Bab(76-109) and BB to Bab(1-89), all his-tagged at the C-terminus. **E.** Growth curves of liquid cultures at 26° of *E. coli* BL21(DE3) pLysS cells expressing different Bab constructs. Bab corresponds to full-length Bab, BASS11 to Bab(76-109) and BB to Bab(1-89) all his-tagged at the C-terminus. IPTG at a final concentration of 0.1 mM was added at time 0. Grey curves are with ITPG, black curves without ITPG. **F.** Colony formation after IPTG induced expression in BL21(DE3) pLysS cells containing a pET24 vector with different Bab-derived constructs (as above) were grown in liquid cultures at 26°C in the presence of 1mM of IPTG for 4 hours and plated on solid 2xYT Amp medium containing no IPTG. Results are given as colony forming units per mL of a culture of an OD_600nm_ of 1.

**Extended data Figure 6.**
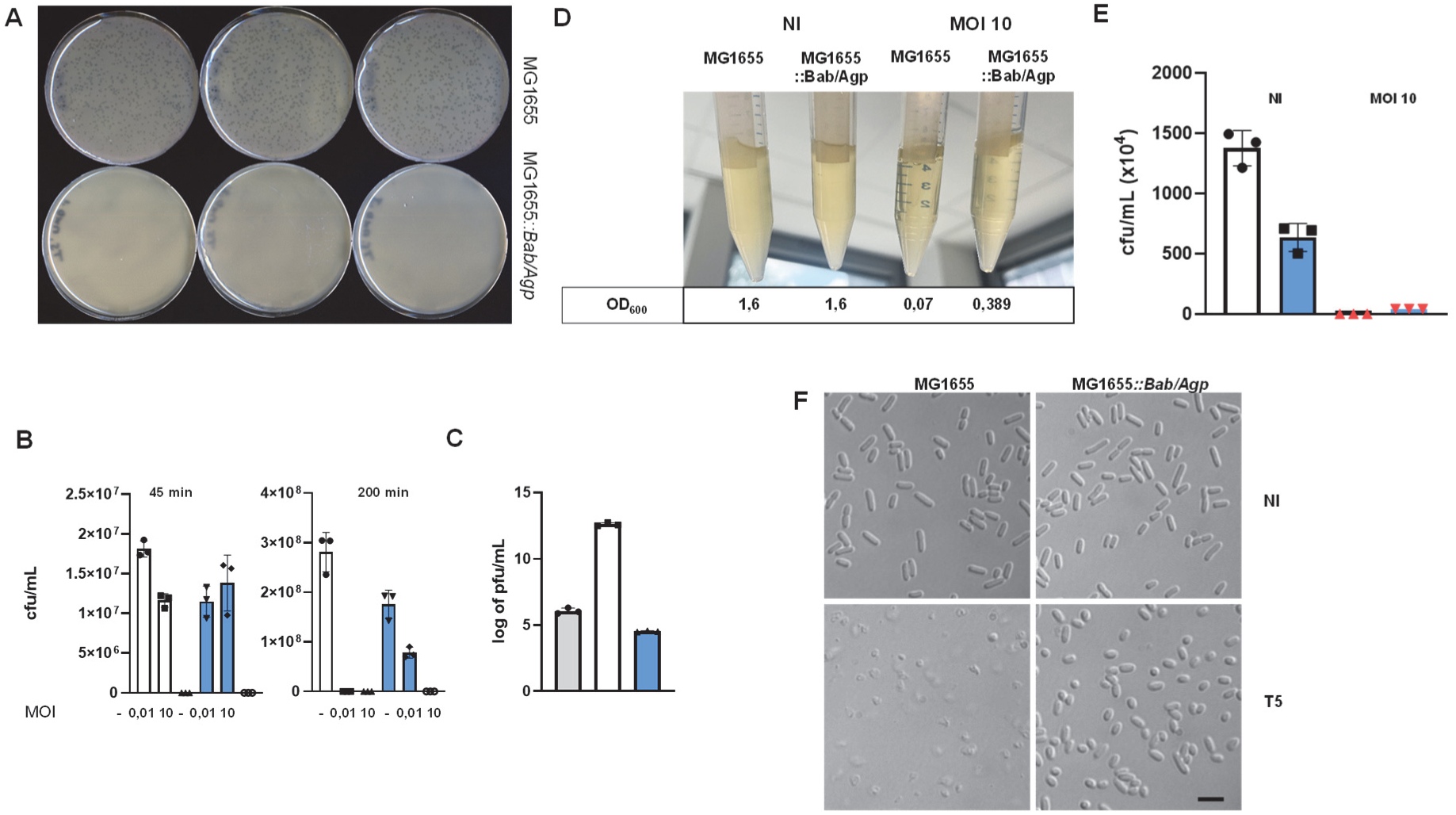
Bab/Agp confers resistance to T5 and induces Abi. **A.** Comparison of plaque formation in MG1655 and MG1655::*Bab/Agp* strains infected with T5. **B.** Counting of colony forming units after infection with T5 of MG1655 (white bars) or MG1655*::Bab/Agp* (blue bars) cells at MOI 0.01 or 10, cultures were infected at OD of 0.1 and plated 45 min or 200 min after infection. **C.** Phage titer in surpernatant of liquid cultures of MG1655 (white bar) or MG1655*::Bab/Agp* (blue bar) cells inoculated with T5 at MOI 0.05 and cultivated for 8 hours at 37°. Cells were infected at OD 0.1. The grey bar corresponds to a control with medium alone inoculated with the same phage titer and incubated similarly for 8 hours at 37°C. The phage titer in the culture supernatant was then measured in spot assays on MG1655 cells. **D.** T5-infected MG1655*::Bab/Agp* cells die but do not lyse. Photographs of cultures of MG1655 and MG1655*::Bab/Agp* cells infected with T5 at MOI 10 (or non-infected, NI) at OD 0.45 and cultivated an additional hour at 37° with agitation. For each culture the measured OD at 600nm is given. **E.** Counting of colony forming units in the cultures shown in E. obtain after plating on LB medium. **F.** Microscopic examination of the cultures shown in D. Scale bar is 4 µm.

**Extended data Figure 7.**
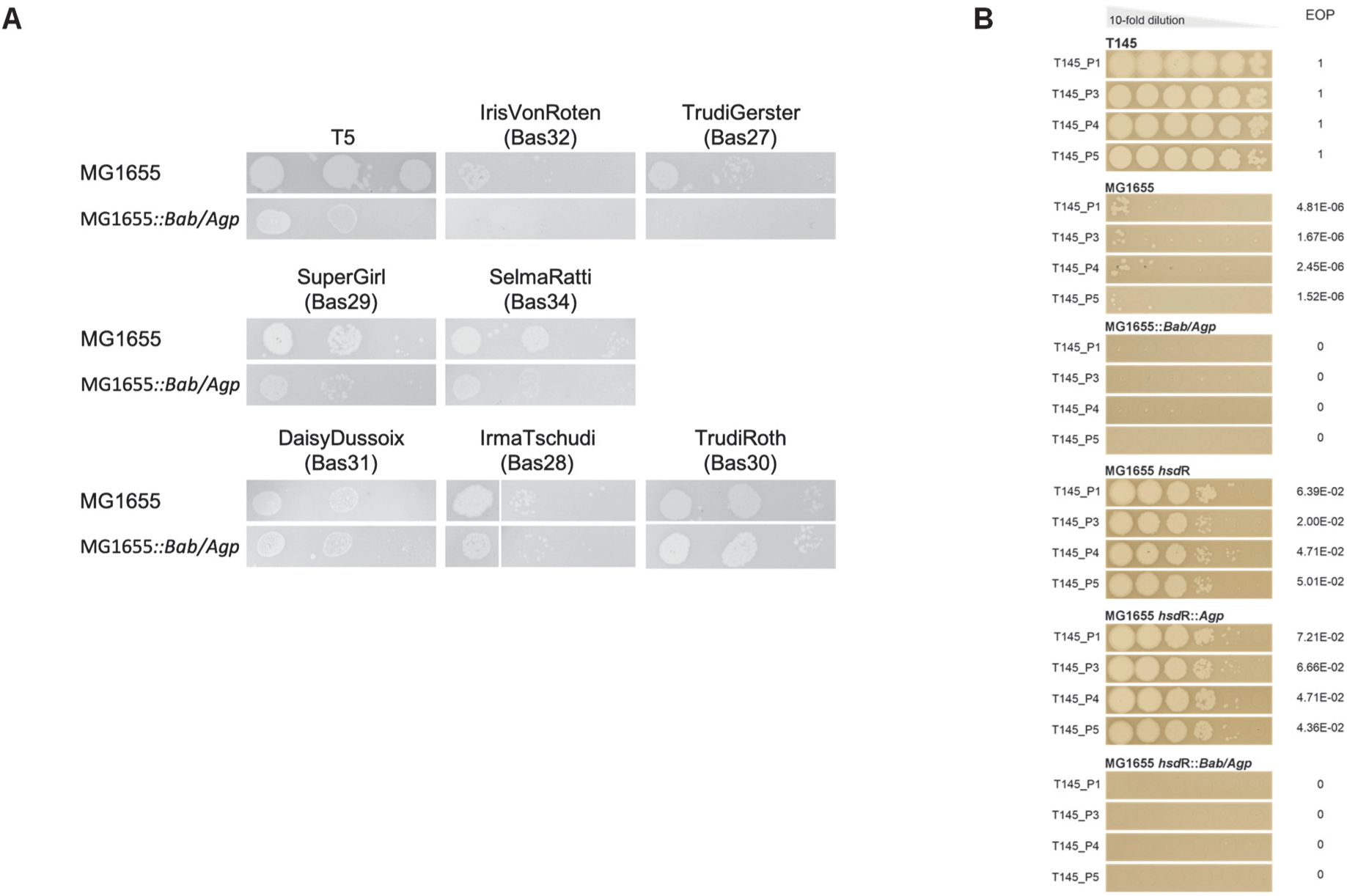
*Bab/Agp* confers resistance across different viral families. **A.** Spot assays with 10-fold serial dilutions of T5-like phages on MG1655 and MG1655::Bab/Agp cells. Phages in the first row exhibit reduction in phage titer on the strain carrying Bab/Agp. In the second row, although no titer reduction is observed, the plaques appear smaller. Finally, the third row displays phages related to T5 which are not affected by the defense system. **B.** Representative pictures of spot assays on the indicated strains of phages T145_P1/3/4/5 that belong to the *Drexlerviridae* family of viruses by contrast to T5 that belongs to the *Demerecviridae* family. The table reports the efficiency of plaquing of these phages on the indicated strains taking as a reference the isolation host, strain T145.

**Extended data Figure 8.**
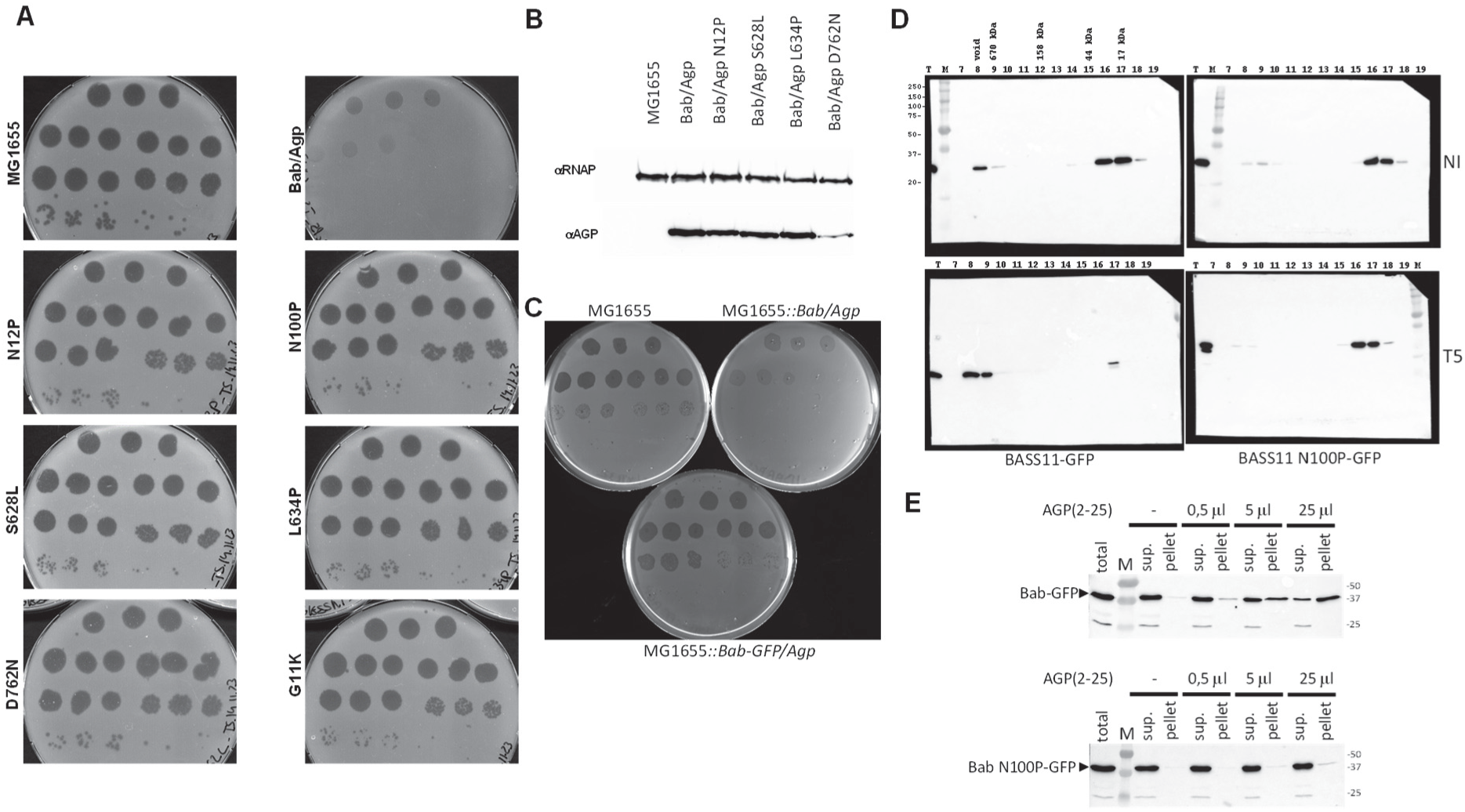
Effect of mutations in Bab and Agp on resistance to T5 and aggregation of BASS11-GFP after T5 infection. **A.** Spot assay of *E. coli* cells of the given genotype infected with 10-fold serial dilutions of T5 in triplicates. Each spot is 4 µL and dilutions are 10^2^ to 10^8^ of the phage stock. **B.** Western-blot of crude extracts of *E. coli* MG1655 cells expressing V5-tagged version of Agp and Agp mutants from an arabinose inducible pBAD vector with an antibody to the V5 tag. As a loading control, the same blot was probed with an antibody to the β’ subunit of *E. coli* RNA polymerase (αRNAP).C. **C.** The GFP tag affects antiphage activity of Bab. Spot assay of *E. coli* cells of the given genotype infected with 10-fold serial dilutions of T5 in triplicates. Each spot is 4 µL and dilutions are 10^2^ to 10^8^ of the phage stock. **D**. Western-blot with anti-GFP antibodies of MG1655 *E. coli* extracts expressing BASS11-GFP or BASS11 N100P-GFP without (NI) or with infection by T5 at MOI 10 fractionated by size exclusion chromatography. Crude extracts were obtained 25 min after infection and fractionated by size exclusion chromatography on a Superdex 200 column, fraction 7 to 19 were analyzed by Western-blot. Position of the SEC molecular weight markers with respect to the analyzed fractions are given. Molecular weight markers of the western-blot (lane M) are given in kDa. Lane T corresponds to the total non-fractionated extract. **E.** Western-blot of *E. coli* MG1655 extracts expressing Bab-GFP or Bab N100P-GFP incubated with various amounts of Agp(2-25) peptide in amyloid conformation and fractionated by centrifugation at 17,000g for 30 min. Agp(2-25) peptide was at 2 mg/mL and was added to 50 µL of crude extract (corresponding to 10^9^ cells). Size of molecular weight markers (lane M) are given in kDa.

**Extended data Figure 9.**
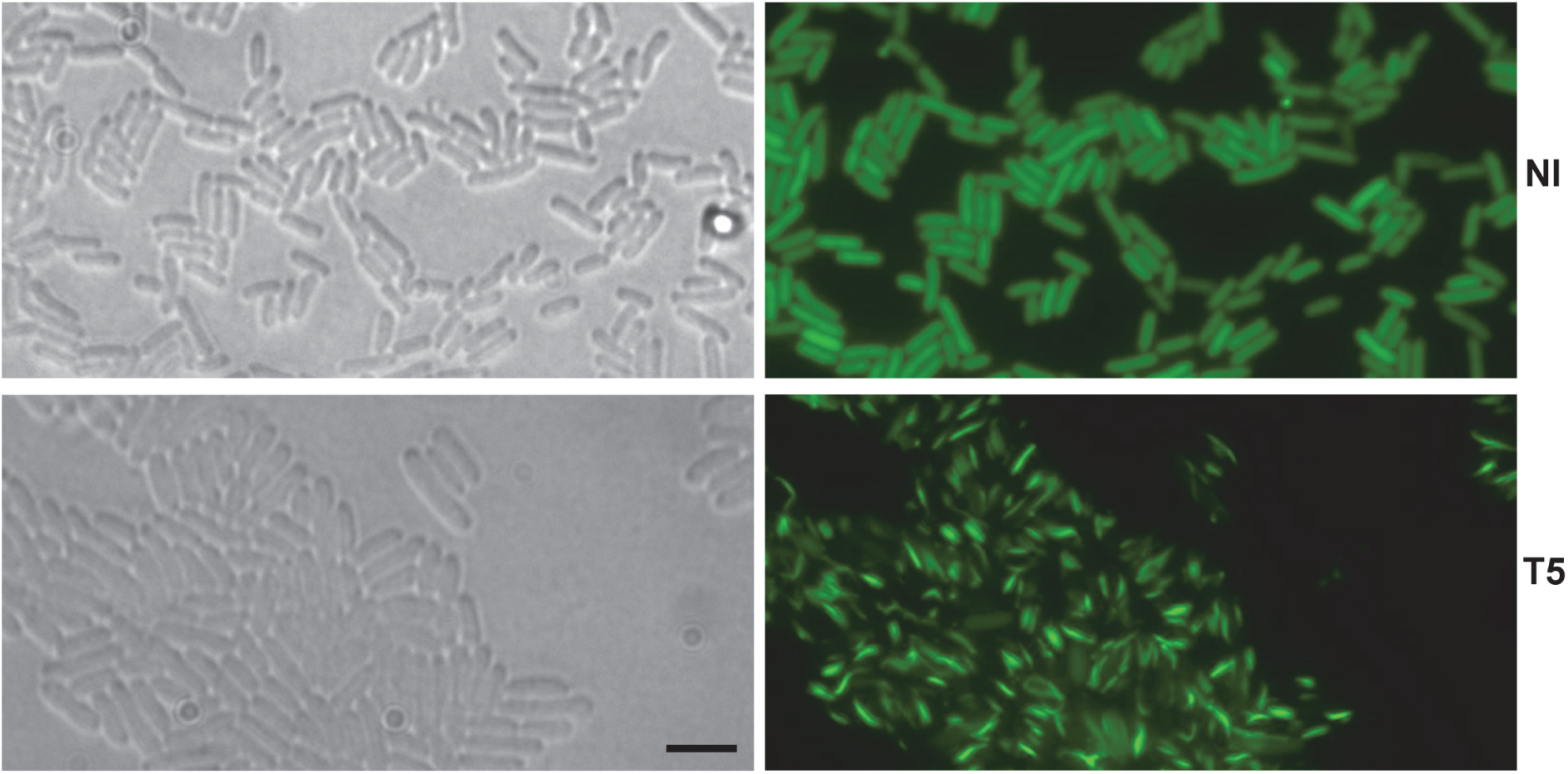
T5 infection leads to formation of BASS11-GFP rods in Bab/Agp cells. MG1655*::Bab/Agp* cells expressing BASS11-GFP were imaged before or 25 min after infection with T5 at MOI 10. Scale bar is 4 µm.

**Extended data Figure 10.**
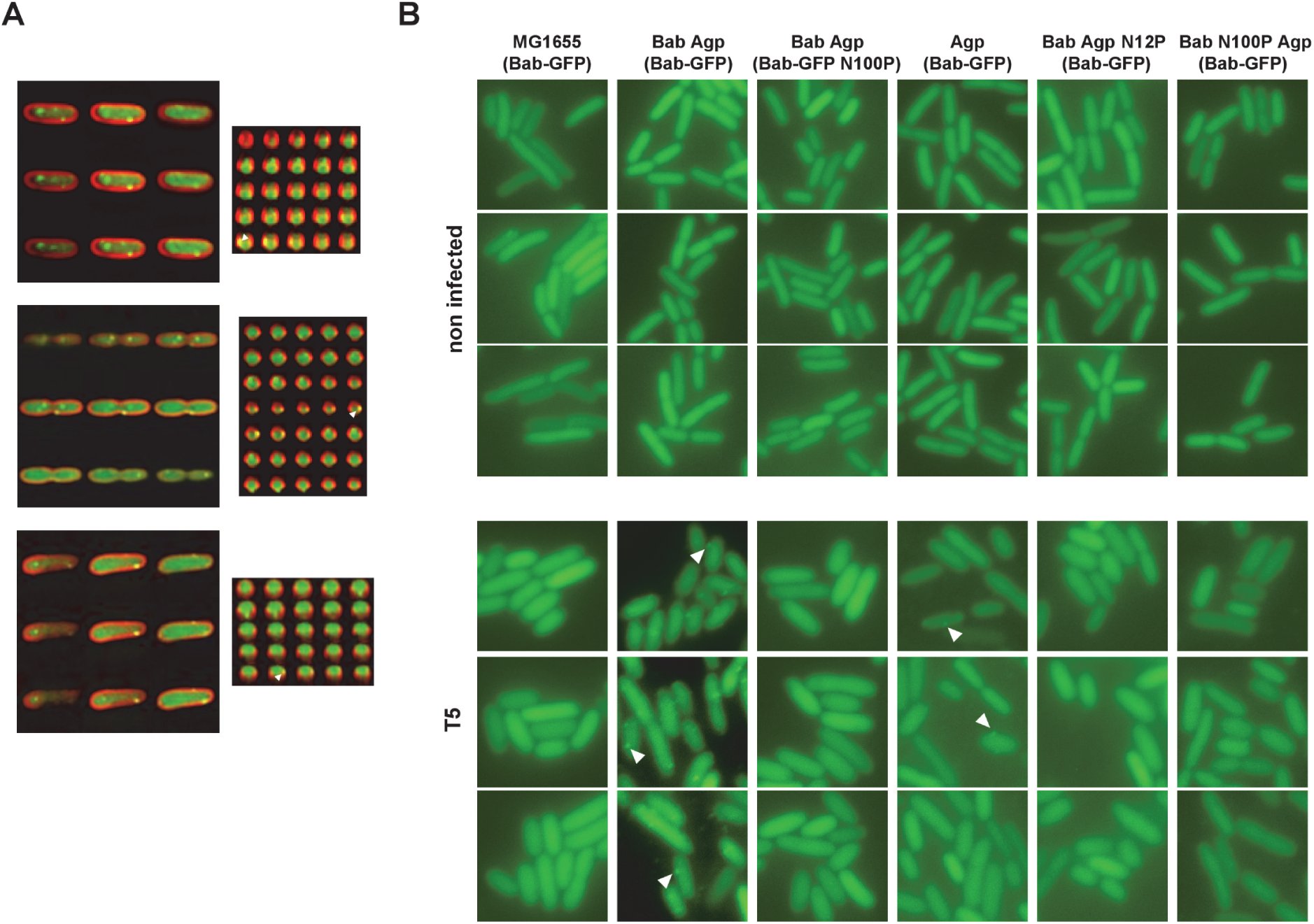
Relocalization of Bab-GFP Membrane alterations after T5 infection. **A.** Representative examples of MG1655*::Bab/Agp* cells expressing Bab-GFP infected with T5 at MOI 10, labelled with Nile Red and imaged 25 min after infection. The imaged gives a series of Z-stacks for Nile red and GFP overlay. On the right, an orthogonal view of the Nile Red/GFP overlay is given and orthogonal views progress from left to right. On each of the orthogonal view a white arrowhead points to an example of a Bab-GFP puncta located close to the cell periphery. **B.** *E. coli* cells expressing Bab-GFP or Bab N100P- GFP and with the given genotype were infected with T5 at MOI 10 and imaged 25 min after infection. A white arrowhead points to examples of Bab-GFP puncta at the cell periphery. Each panel is 10 x 10 µm.

**Extended data Figure 11.**
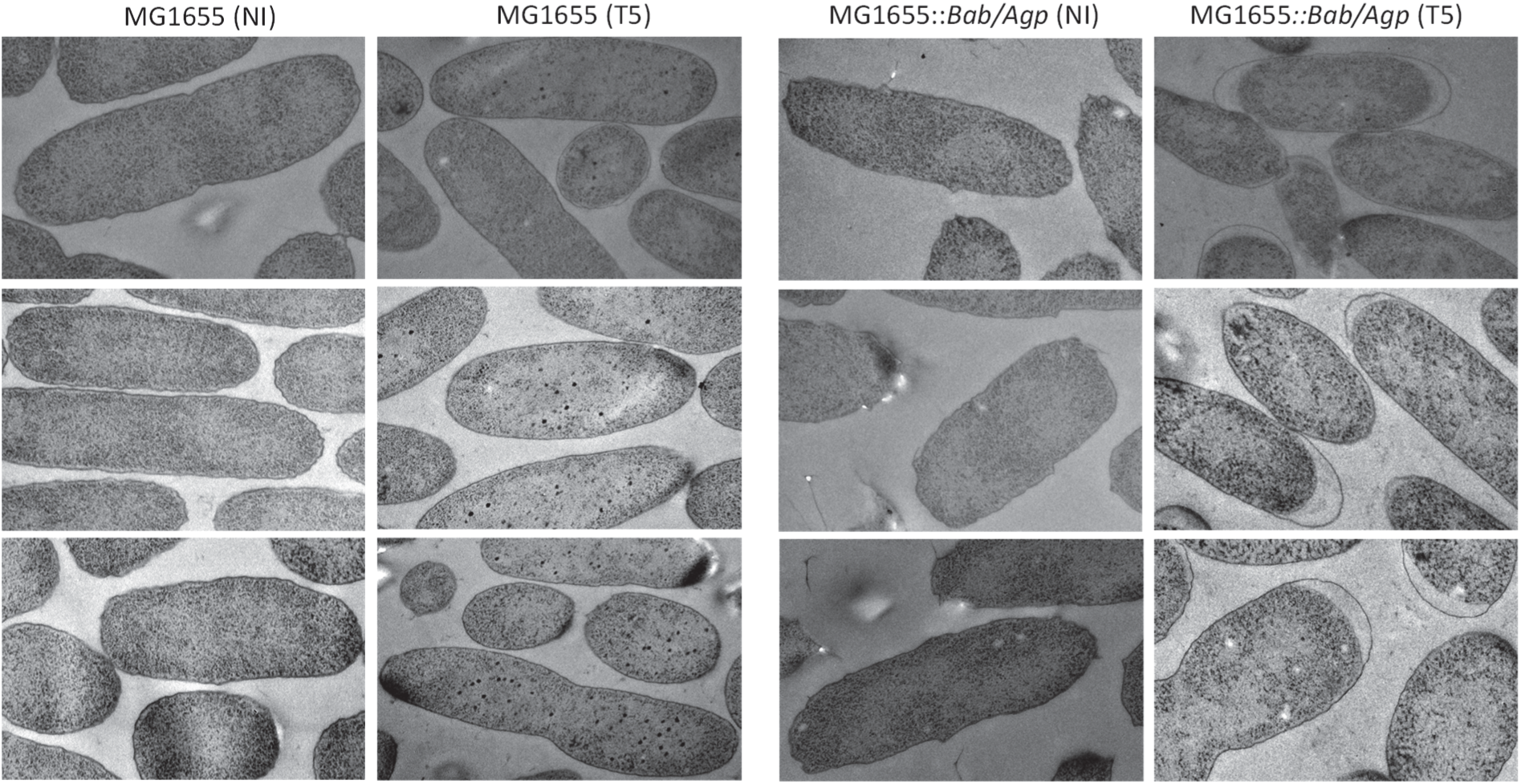
Membrane alterations after T5 infection. Transmission electron microscopy of MG1655 and MG1655*::Bab/Agp* cells were infected with T5 at MOI 10 (T5) or non infected (NI). Cells were harvested for fixation 25 min after infection. Each panel is 2 x 3 µm.

**Extended data Figure 12.**
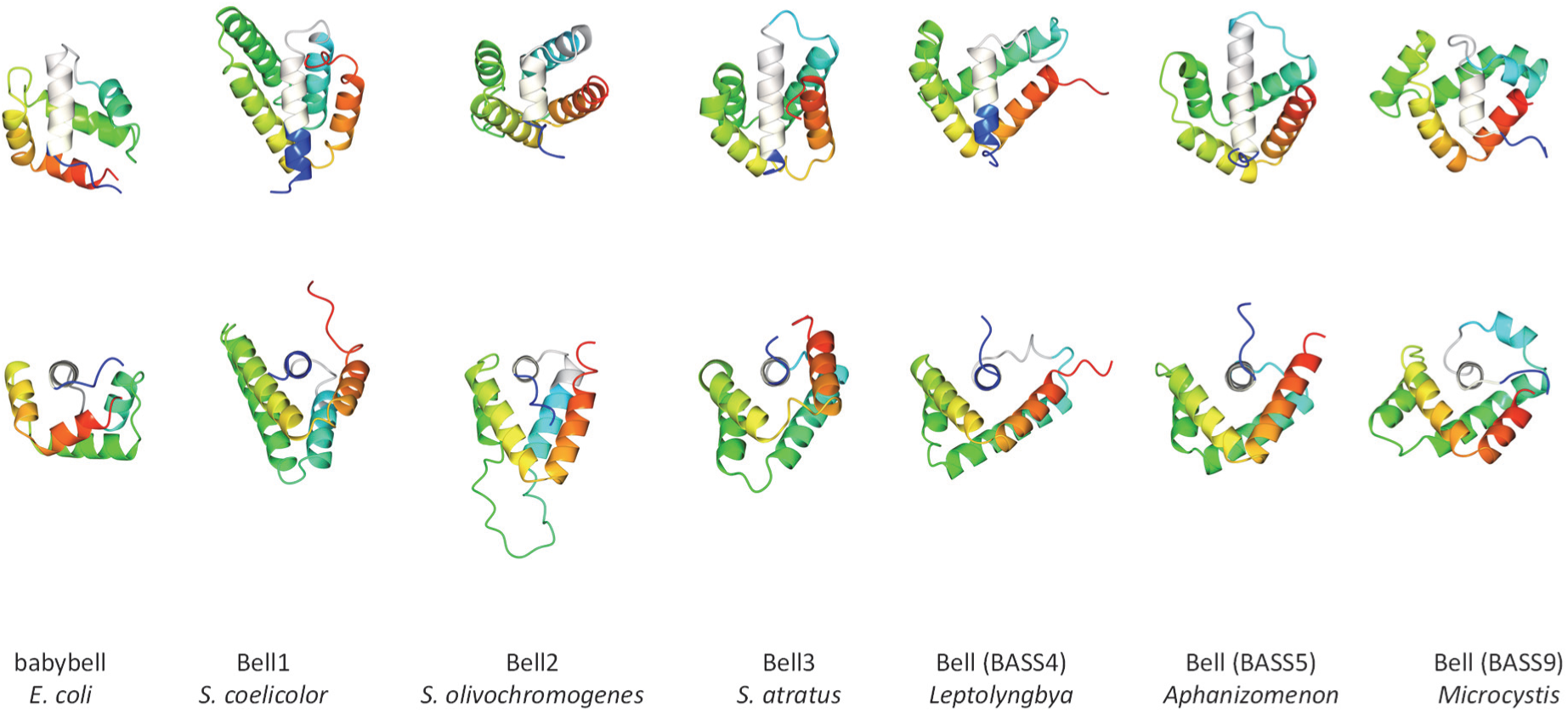
Position of the conserved MEPIS-motif region in Bab and related bacterial proteins. The experimental Bab structure and the predicted AlphaFold structures of the Bell domain belonging to different clusters (as given in extended data Fig. 1D) with a rainbow coloring with blue to red from N- to C-terminus. The predicted pore-lining helix (as predicted with MEMSAT-SVM) is given in beige in each case. Bell1, 2 and 3 are from *Streptomyces* species (actinobacteria) while the three Bell domains on the left, associated to the BASS4, 5 and 9 motifs are from cyanobacterial species.

**Extended data Figure 13.**
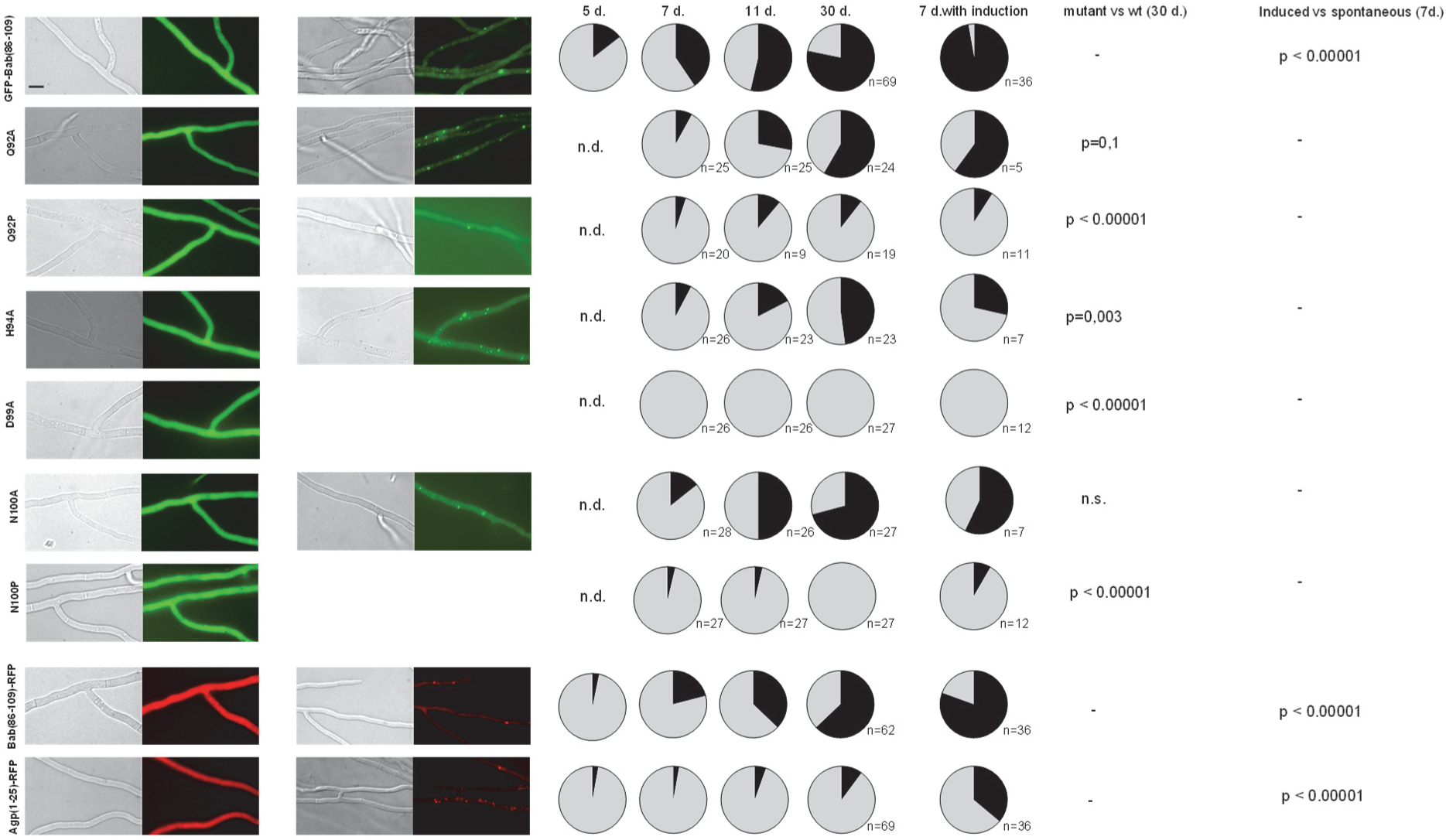
GFP-BASS11 forms foci *in vivo* when expressed in *P. anserina.* **A.** representative examples *P. anserina* transformants displaying the diffuse fluorescence state ([b*] state) and foci state ([b] state) of GFP-Bab(86-109), Bab(86-109)-RFP and Agp(1-25)-RFP and of mutant versions of GFP-Bab(86-109). No images for the foci state are given for D99A and N100P because the dot state is almost never observed. Scale bar is 5 µm. **B.** The pie charts give the fraction of transformants showing diffuse fluorescence (grey sector, [b*] state) or foci (black sector, [b] state), 5, 7, 11 and 30 days after transformation, (n.d., not determined). In each case, the number of transformants that were analyzed is given (n). Transformants were also confronted with strains showing foci (7 d. with induction) and the fraction of strains showing foci after this confrontation is given. P-values (two-tailed Fischer tests) for the foci formation at 30 days in mutants compared to wild-type are given as well as for spontaneous compared to induced foci formation (n.s., not significant).

**Extended data Figure 14.**
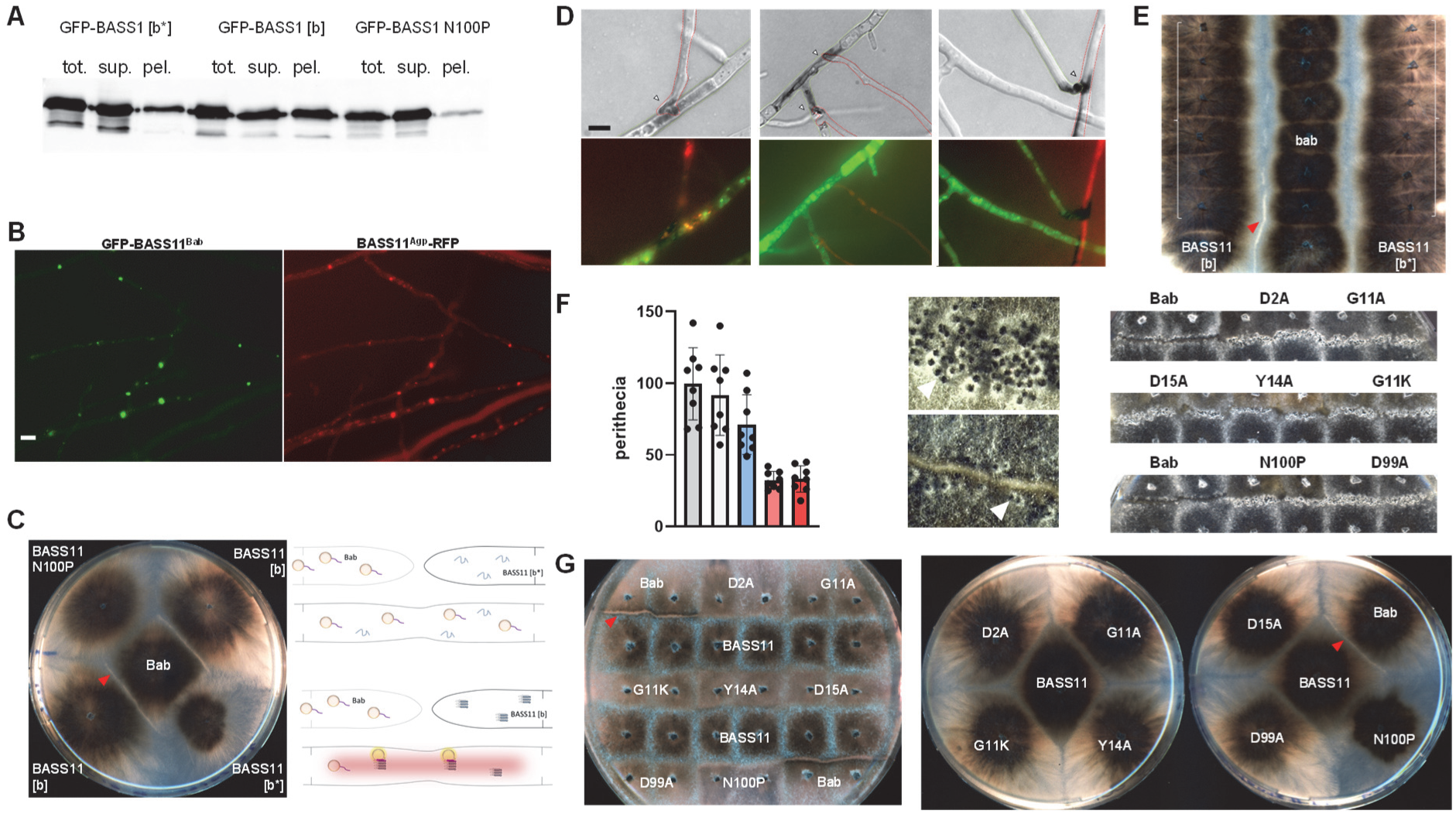
BASS11 aggregate formation in *P. anserina* and Bab/BASS11 incompatibility. **A.** Western-blot of *P. anserina* crude extracts expressing GFP-BASS11 (GFP-Bab(86-109)) in the [b*] and [b] state or GFP-BASS11 N100P fractionated by centrifugation at 17,000g probed with an anti-GFP antibody. **B.** Colocalization of GFP-BASS11^Bab^ (GFP-Bab(86- 109)) and BASS11^Agp^-RFP (Agp(2-25)-RFP) foci in *P. anserina*. Scale bar is 5 µm. **C.** Barrage reaction between a strain expressing Bab-GFP (Bab) and strains expressing GFP-BASS11 is soluble [b*] and aggregated [b] state on corn meal agar medium. The red arrowhead points to the barrage reaction. On the right an interpretative model of Bab/[b] incompatibility is given. In this model, in [b*] strains BASS11 is soluble, upon cell fusion with a strain expressing Bab, no activation of Bab occurs and the fusion cell is viable. In [b] strains, BASS11 is aggregated into an amyloid state and upon fusion with a strain expressing Bab, Bab is converted and activated and causes death of the fusion cell. **D.** Cell death in fusion cells of strains expressing Bab (Bab(86-109)-RFP) and BASS11 (GFP-Bab(86-109)). The upper panels corresponds to methylene blue staining for dead cells, the lower panel to a GFP/RFP overlay. Arrowheads point to the presumed fusion site. **E.** GFP-BASS11 [b] prion propagation assay based on acquisition of the barrage reaction to a tester strain expressing Bab. Two rows of six [b*] state strains expressing GFP-BASS11 were inoculated on the plate (grey bracket) together with a row of strains expressing Bab (center row). The left and right row differ by the strain inoculated at the bottom of the plate, either [b*] (bottom right) or [b] strain (bottom left). Note the barrage formation in the confrontation zone in the left row, indicating that [b*] strains have been converted to the [b] state at the time of contact with the Bab-tester row. **F.** Sexual incompatibility associated to the Bab/BASS11 interaction. Counts correspond to the number of fertilized sexual organs (perithecia) formed in the following crosses : *ΔhellpΔhet-sΔhellf::Bab* x *ΔhellpΔhet-sΔhellf* (*ΔΔΔ::Bab x ΔΔΔ)* (shades of grey, two independent crosses), *ΔΔΔ::GFP- BASS11* x *ΔΔΔ* (blue, one cross) and *ΔΔΔ::Bab* x *ΔΔΔ::GFP-BASS11* (red and pink, two independent crosses) crosses. Counts are numbers of perithecia per cm in the confrontation zone. Images of the confrontation zone showing individual perithecia (white arrowheads) are given with the upper image is *ΔΔΔ::Bab x ΔΔΔ* and the lower image, *ΔΔΔ::Bab* x *ΔΔΔ::GFP-BASS11*. On the right stripes of confrontation zones show that mutations in Bab abolish sexual incompatibility. In the right panel, strains expressing Bab and Bab mutants as marked were crossed with strains expressing aggregated GFP-BASS11 (*ΔΔΔ::Bab* x *ΔΔΔ::GFP-BASS11*). All tested mutations increase the number of fertilized sexual organs (perithecia). **G.** Mutations in the BASS11-motif and the N-terminal region of Bab abolish vegetative incompatibility. Mutations in Bab abolish the barrage reaction in confrontations with strains expressing aggregated BASS11. The confrontations are shown on synthetic medium (left) and corn meal agar (right). The red arrowhead points to the barrage reaction between the strain expressing Bab and GFP-BASS11.

**Extended data Figure 15.**
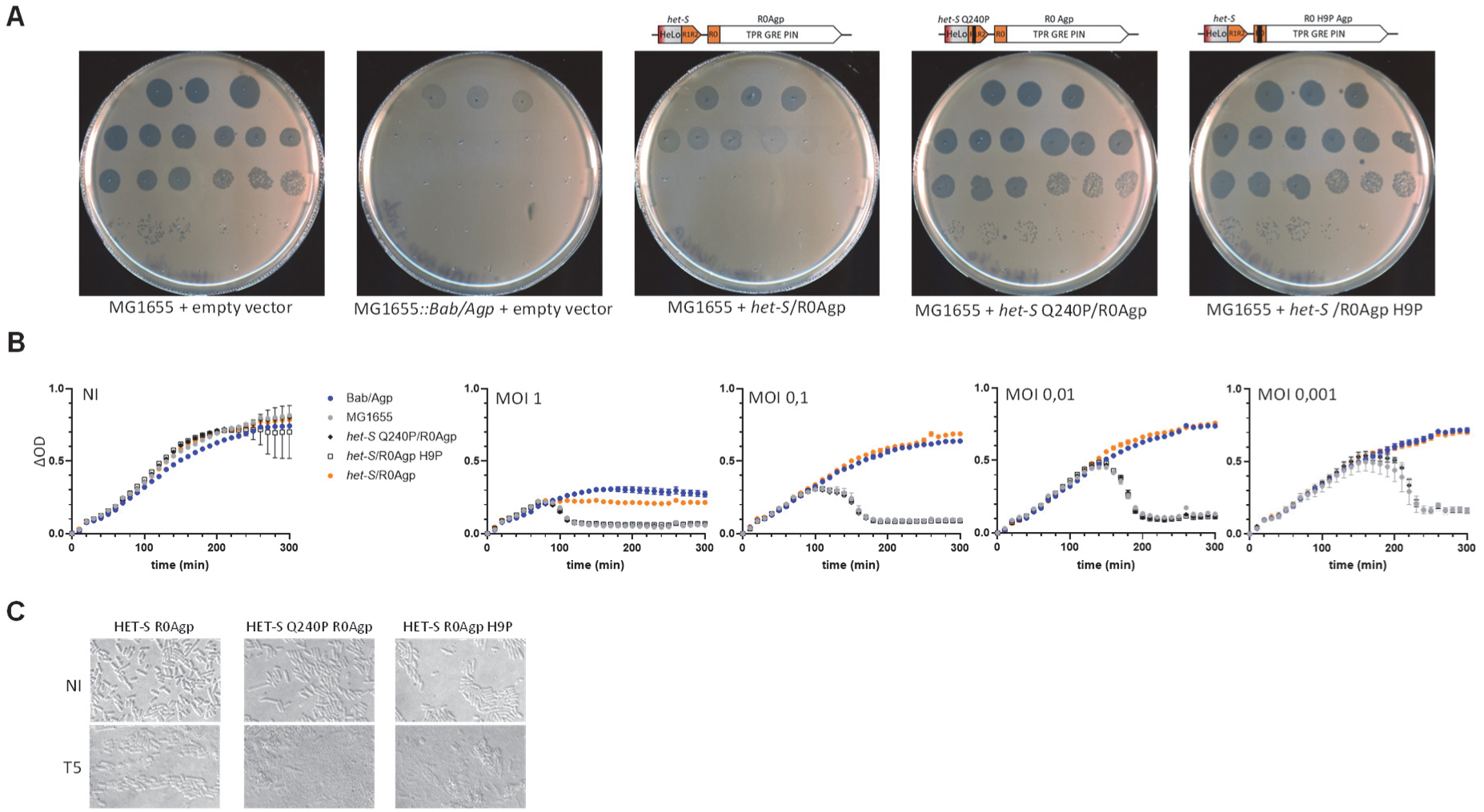
*het-S*/R0Agp confers resistance to T5. A Spot assay with 10-fold serial dilutions of T5 on lawns of MG1655 cells and containing a plasmid with *het-S/R0Agp* and mutant version of the same construct *(het-S* Q240P and *R0Agp* H9P). MG1655 cells and MG1655*::Bab/Agp* cells carrying the empty vector are used as controls. Each dilution is spotted in triplicates, serial dilutions are from top to bottom and left to right. **B.** Growth curves of strains used in A. and infected with T5 at different MOIs as indicated.Cells of strains expressing HET-S/R°Agp, HET-S Q240P/R0Agp and HET-S/R0Agp H9P infected with T5 at MOI 10 and imaged 45 min after infection, (images are 45x30 µm). Cell ghost are present in HET-S R0Agp cultures while most cells are lysed in the Q240P and H9P mutants. Cells detected in HET-S/R0Agp are dead based on colony forming ability (<10^-3^ cfu/cell).

## Supplementary Tables

Table S1. Genbank accession for sequences listed in extended data Fig.1A

Table S2. ArchCandy and AmyloComp scores of selected BASS-motifs

Table S3. List of chromosomal insertions

Table S4. Primers

Table S5. Plasmid constructs

Table S6. Diffraction data collection statistics for Bab(1-90) C2 crystals measured at the copper Kα wavelength

